# The early communication stages between serine proteases and enterovirus capsids in the race for viral disintegration

**DOI:** 10.1101/2023.08.30.555488

**Authors:** Marie-Hélène Corre, Benjamin Rey, Shannon C. David, Shotaro Torii, Diego Chiappe, Tamar Kohn

**Affiliations:** Laboratory of Environmental Chemistry, Environmental Engineering Institute (IIE), School of Architecture, Civil and Environmental Engineering (ENAC), Ecole Polytechnique Fédérale de Lausanne (EPFL), 1015-CH Lausanne, Switzerland.; Proteomics Core Facility, School of Life Sciences, Ecole Polytechnique Fédérale de Lausanne (EPFL), 1015-CH Lausanne, Switzerland.

## Abstract

Enteroviruses are human pathogens known to challenge water safety^1,2^. Among the microbial stressors found in water, bacterial serine proteases contribute to the control of enterovirus persistence^3^. However, the structural interactions accounting for the susceptibility of enteroviruses to proteases remains unexplained. Here, we describe the molecular mechanisms involved in the recruitment of serine proteases to viral capsids. Among the virus types used, coxsackievirus A9 (CVA9), but not CVB5 and echovirus 11 (E11), was inactivated by Subtilisin A in a host-independent manner, while Bovine Pancreatic Trypsin (BPT) only reduced CVA9 infectivity in a host-dependent manner. Predictive interaction models of each protease with capsid protomers indicate the main targets as internal disordered protein (IDP) segments exposed either on the 5-fold vertex (DE loop VP1) or at the 5/2-fold intersection (C-terminal end VP1) of viral capsids. We further show that a functional binding protease/capsid depends on both the strength and the evolution over time of protease-VP1 complexes, and lastly on the local adaptation of proteases on surrounding viral regions. Finally, we identified three residues on CVA9 capsid that trigger cleavage by Subtilisin A, one of which acts as a sensor residue contributing to enzyme recognition on the DE loop. Overall, this study describes an important biological mechanism involved in enteroviruses biocontrol.

## Introduction

Serine proteases are important environmental contributors of enterovirus biocontrol^3^. However, not all virus types are equally susceptible to these enzymes and the range of specificity of serine proteases does not solely explain virus inactivation^4^. Accordingly, we performed a structural and functional study of the interaction between enterovirus capsids (CVA9, CVB5 and E11) and serine proteases (Subtilisin A from *Bacillus licheniformis*, Trypsin from bovine pancreas (BPT)), to unravel the molecular steps leading to viral inactivation.

As first partner of interaction, *Enterovirus* capsids are giant ∼5’500-kilodalton (KDa) rounded substrates, composed of 60 repeating units of the four viral proteins VP1, VP2, VP3 (protomer) and VP4 (Extended data Fig. 1 a)^5–7^. The overall capsid architecture is built by the association of twelve repeats of five protomers (12 pentamers), tightly bound together to form an icosahedral shell with a 5:3:2 rotational symmetry (Extended data Fig. 1 a, b) ^5,6,8^. From this structural arrangement, VP1, VP2, VP3 cover the outer lattice and share a similar jelly roll β-sandwich fold, while the minor protein VP4 is plated to the inner surface of the capsid (Extended data Fig. 1 a-e)^5,6,9^. The eight β-sheets of each jelly roll fold, conventionally designated by the letters B to I, are linked to each other by internal disordered protein (IDP) loops, some of them contributing greatly to the plasticity of both the 5-fold vertex (‘mesa’) and the 3-fold proper-like protrusion of the capsid (Extended data Fig. 1 d, e)^10,11^. While *Enterovirus* VP1s share the same core structure, their sequences are known to differ from 279 to 317 amino acids in length depending on virus types^12^. As a result, the C-terminal end of some VP1s (∼5 to 20-mer length) is exposed around the 5/2-fold axes intersection of the capsid and constitutes an additional structurally accessible IDP segment^13^.

**Figure 1:**
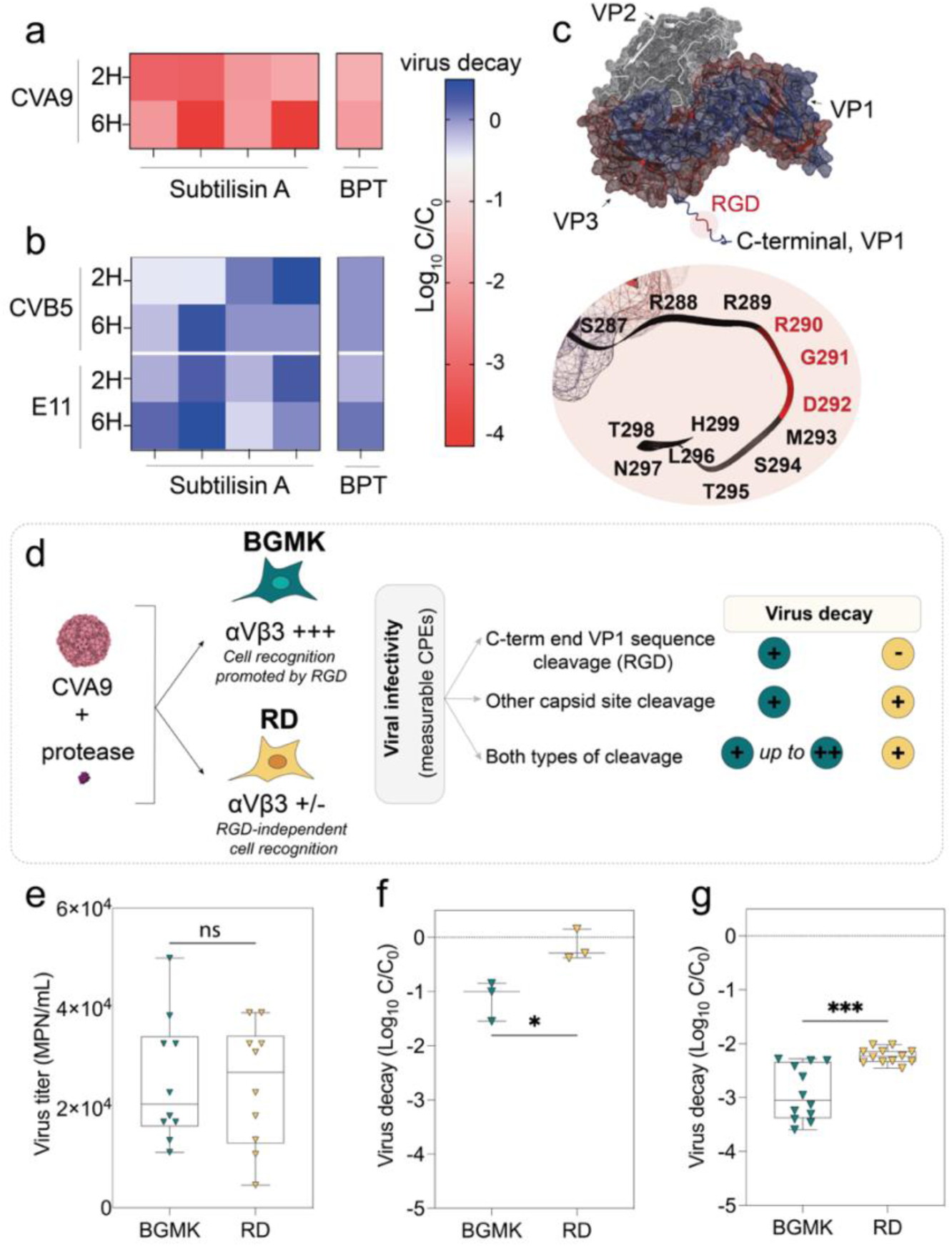
Enterovirus inactivation by serine proteases is both virus type and protease specific. **a-b.** Virus decay (Log_10_ C/C_0_) of **(a)** CVA9 or **(b)** CVB5 and E11 after an exposure to serine proteases. Each virus has been exposed to 20 μg/mL of Subtilisin A (n=4) or BPT (n=1) for 2 or 6 hours. Infectivity experiments have been performed on BGMK cells and cytopathic effects (CPEs) measurements have been done 4-days post-infection. **c.** Visualization of the CVA9 C-terminal end VP1 segment (strain Griggs, 1d4m), carrying a RGD motif. Modeling of CVA9 VP1 anchoring the C-terminal end of the protein has been performed using Modeller. **d.** Screening procedure used to demonstrate the RGD-independent viral decay of CVA9 by serine proteases. **e.** Comparison of CVA9 viral titers 5-days post-infection on BGMK and RD cells (*t*=0.1667, *p*-value:0.8695 (ns); two-tailed unpaired *t*-test, n=10). **f-g.** Comparison of CVA9 viral decay measured on BGMK and RD cells after an exposure either with 20 ug/mL BPT **(f)** *(t*=3.583, *p*-value:0.0256 (*); two-tailed unpaired *t*-test, n=3) or Subtilisin A **(g)** (*t*=4.614, *p*-value:0.0005 (***); two-tailed unpaired *t*-test, n=12) for 2 hours. For each experiment involving a treatment with proteases, the LoD of the assay was 5-log_10_. For all negative control experiments of inactivation (C0), PBS has been used to replace serine proteases. **e-g**. For each box plot, the horizontal bar within the interquartile range corresponds to the median of values. The vertical bar of each dataset corresponds to the distribution of minimum and maximum values.

As second partner of interaction, serine proteases are ∼25 KDa monomeric enzymes described to catalyze the hydrolysis reaction of peptide bonds under the action of a nucleophilic serine^14^.

Trypsins and subtilisins are among the most studied serine proteases and are known to share a similar catalytic pocket geometry^15–19^. The catalytic site of such enzymes is formed by a triad folded structure, which ensures a charge relay between an aspartic acid, a histidine, and the nucleophilic serine^15,18,20^. Moreover, an adjacent residue (glycine or asparagine) is hydrogen bonded to the nucleophilic serine, forming an oxyanion hole used to stabilize the reaction intermediate^21,22^. Serine proteases are however distinguished from each other by the specificity of substrate they can hydrolyze. While trypsins specifically hydrolyze the scissile bond following basic amino acids (K, R)^23,24^, subtilisins deploy a much broader spectrum of action and tend to include most amino acids as potential substrate, with the exception of asparagine and tryptophane^25–29^.

## Results

### Virus type-specific sensitivity to serine proteases

To investigate the specificity of interaction between capsids and serine proteases leading to virus inactivation, we first selected sensitive and insensitive virus types to Subtilisin A and BPT. As previously described, CVA9 is sensitive to several proteases including BPT and a subtilisin^4^. We therefore selected this virus type to set up our antiviral assay with the two proteases. To this end, an environmental CVA9 isolate was exposed for 2 or 6 hours to 20 μg/mL of each protease, and the virus’ ability to infect Buffalo Green Monkey Kidney (BGMK) cells was measured before and after exposure. The cytopathic effects (CPEs) measured on cells after an exposure with each protease were used to demonstrate the reduction of viral infectivity (Log_10_ C/C_0_) (Fig.1 a). Independent of the treatment time, exposure to Subtilisin A led to 1.5 to 3.5-log_10_ reduction of CVA9 infectivity, whereas a similar exposure with BPT reduced viral infectivity by 1.5-log_10_. With the aim of finding protease-resistant viruses, we reproduced this experiment with the reference strains CVB5 (strain Faulkner) and E11 (strain Gregory), for which sensitivity to these two proteases was unknown. For all experimental conditions, no reduction of the infectivity was observed (Fig.1 b), indicating that the capsid integrity of these two virus types is preserved upon exposure with both proteases.

### Protease-specific inactivation of CVA9

Unlike CVB5 and E11, CVA9 exposes on its capsid outer surface a 18-mer hyperflexible C-terminal VP1 sequence (IDP) carrying an RGD motif (Fig.1 c)^13,30,31^. This motif is described to promote CVA9 recognition to BGMK cells through the integrin cell signaling pathway (αVβ3, αVβ6) during the early stages of infection^32–34^. This C-terminal sequence is also known to be a valuable substrate for some proteases, though its cleavage does not influence CVA9 infection via an integrin-independent cellular bypass process^35,36^. To investigate whether Subtilisin A and BPT attack this IDP on CVA9 capsid, we developed a differential cell culture approach based on integrins recognition (Fig.1 d). We exposed CVA9 to each protease, before measuring its infectivity on both BGMK cells (RGD-promoted cell recognition) and human Rhabdomyosarcoma (RD) cells, known to be permissive to CVA9 in an RGD-independent manner^36,37^. Prior to this experiment, the percentage of cells expressing each integrin type (*i.e*., αVβ3, αVβ6) was assessed by flow cytometry for both cell lines (Extended Data Fig. 2 a, b). We counted on average 9.5-fold more αVβ3-type integrin on the surface of BGMK cells (40-90% positive cells) than on RD cells (1-6% positive cells), whereas αVβ6-type integrin expression was similar for both cell lines (5% positive cells). We then monitored the CPEs induced by CVA9 on RD and BGMK cells to standardize our viral assay and to further study the C-terminal end sequence cleavage by serine proteases (Extended Data Fig. 2 c). Similar viral titers were measured on both cell lines 5-days post-infection (∼ 0.5.10^5^ MPN/mL, *p*-value = 0.8695, unpaired *t*-test) (Fig. 1 e), confirming that neither random nor cell-line-specific factors (*i.e.* integrins proportion, other attachment proteins, protein from media) influenced the infectivity of CVA9 on either cell lines. We then applied the same protocol after an exposure of CVA9 virions to 20 μg/mL of BPT, known to only cleave the C-terminal end VP1 sequence for this virus type^35^. We measured, 5-days post-infection, a reduction (1-log_10_) of CVA9 infectivity on BGMK cells but not on RD cells (n=3, *p*-value = 0.0116, unpaired *t*-test), supporting a differential infection route of the two cell lines after the C-terminal end sequence cleavage (Fig. 1 f). In contrast, exposure to Subtilisin A led to a reduction of infectivity on RD cells (2-log_10_), suggesting that CVA9 is inactivated in a RGD-independent manner by this protease (Fig. 1 g). This inactivation was more extensive when measured on BGMK cells (3.5-log_10_), consistent with an additional cleavage of the C-terminal end sequence by Subtilisin A (n=12, *p*-value = 0.0003, unpaired *t*-test with Welch’s correction). While the C-terminal sequence exposed on the capsid surface of CVA9 is thus easily cleaved by both proteases, these data strongly suggest that Subtilisin A also acts on another part of the capsid of this virus type, thereby leading to virus inactivation.

### Multimeric folding of capsid protomers

To further study the interaction of CVA9 with serine proteases at the molecular level, we first sought to model the smallest part of the viral capsid that accounts for the structural constraints brought by the icosahedron assembly. Our rationale led us to consider the modeling of a capsid protomer of CVA9 (heteromer: VP1, VP2, VP3) using AlphaFold2-multimer (AF2-M)^38,39^. Since the CVA9 strain used in this study was an environmental isolate, we first confirmed the amino acid conservation of all purified VPs (Extended Data Fig. 3 a, b) with those from the reference strain Griggs (PDB:1d4m) using mass spectrometry. Based on the sequence coverage achieved after each protein digest, we found 97.7% of VPs amino acids conservation between the environmental isolate and the reference strain (Extended Data Fig. 3 c-e). Due to the high sequence conservation between the environmental isolate and the reference strain, we therefore used the primary sequences of CVA9 Griggs as input for the folding process. The overlay analysis of the top one rank modeled structure on the experimental structure (PDB:1d4m) estimated an overall Root Meaning Square Deviation (RMSD) value of 2.85 Å and a model confidence of 85.2, reflecting a good quality of folding (Extended data Fig. 4 a). Moreover, the representation of the protomer as a function of the predicted local Distance Difference Test (plDDT) achieved for each atom, led to identify eight viral IDPs (7 loops, C-terminal end VP1) folded with lower confidence (plDDT value < 70), which are exposed on the external surface of the capsid (Fig. 2 a, Extended data Fig. 4 b-f). Finally, the representation of the same protomer as a function of the Predicted Aligned Error (PAE) indicate a strong confidence of the position for each VP in the folded structure, though the position of N/C-terminal end segments (all VPs) and one loop (DE, VP1) remained imprecise (Fig. 2 b). With the goal to use CVB5 and E11 as negative controls for molecular modeling experiments with serine proteases, we also modeled the two protomers of CVB5 strain Faulkner (PDB:7c9y) and E11 strain Gregory (PDB:1h8t) using the same methodology (Extended data Fig. 4 g-h). The modeled structures obtained for each viral type were also predicted with high confidence (CVB5: 92.2 (rank 1), E11: 90.3 (rank 1)), though small differences in the confidence level associated with specific loops stand out between the two. For E11, the analysis also integrated a dipeptide at the C-terminal end of VP1, which was missing from the experimental structure (1h8t) and confirm the exposure of an unstructured segment (11-mer peptide) on the capsid surface.

**Figure 2:**
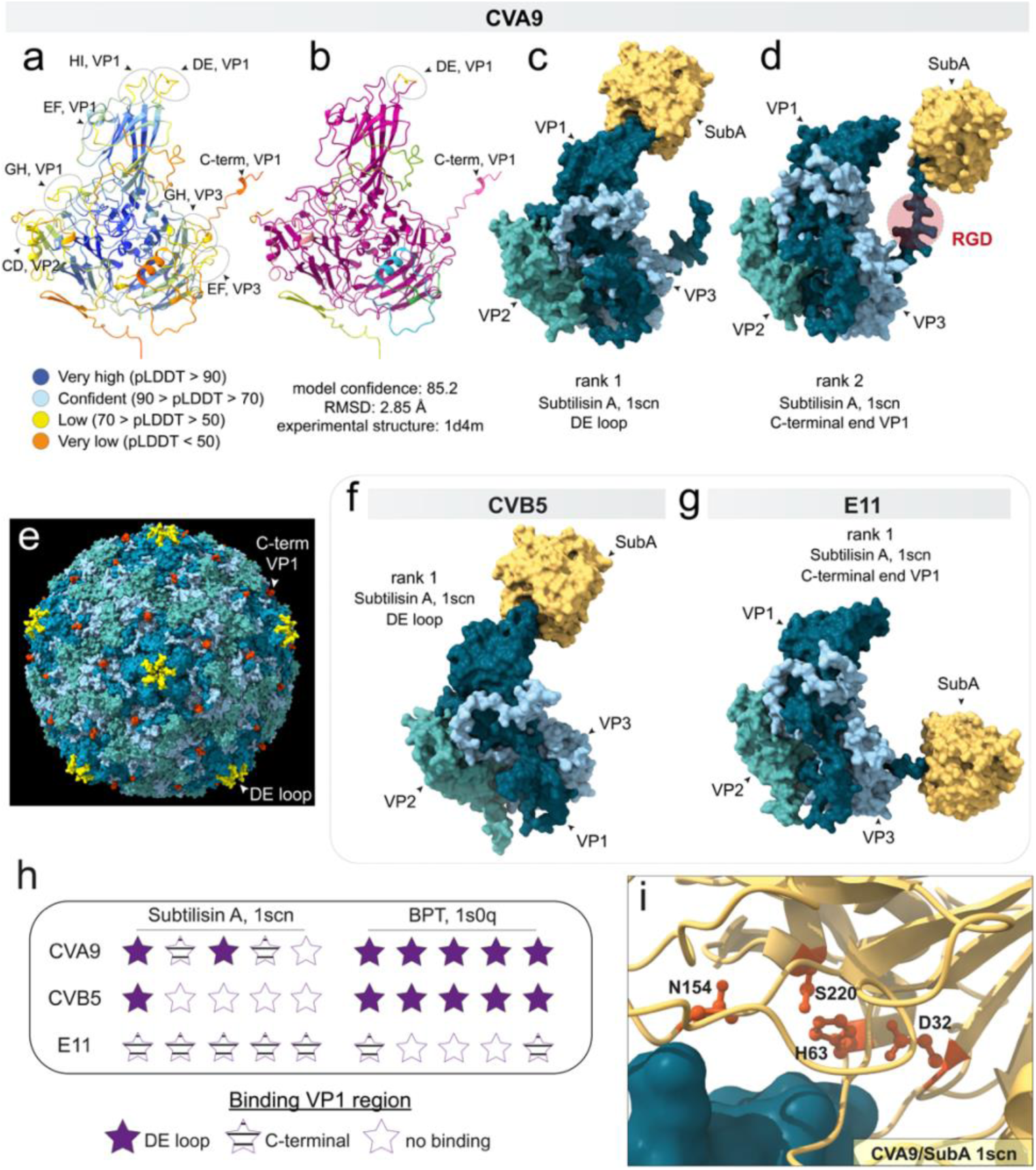
Predictive binding of serine proteases on two viral IDPs. **a-b.** Ribbon diagram of CVA9 protomer modeled by AF2-M (rank 1), displaying **(a)** the confidence level of the model colored by plDDT or **(b)** the confidence according to PAE score matrix. On **a,** the circled areas correspond to jelly roll fold loops predicted with low confidence and found at least partially on the external side of the capsid shell. On **b**, purple ribbons correspond to the residue positions well predicted by the analysis. The other colors correspond to DE loop (VP1), N-terminal VPs and C-terminal VP1, for which residue positions remain unconfirmed by the folding procedure. **c-d.** Surface view of the two models of interaction between Subtilisin A and CVA9 protomer returned by AF2-M. **e.** Surface view of CVA9 full capsid (1d4m) displaying in yellow the joining of five VP1 DE loops on the symmetry center of the 5-fold axis (vertex), and in light red the last residue resolved by X-ray diffraction, preceding the external VP1 C-terminal end segment**. f-g.** Surface view of the models (rank 1) between Subtilisin A and **(d)** CVB5 or **(g)** E11 returned by AF2-M. **h.** Overview of the predicted protein interfaces between each viral protomer and the two proteases of this study. **i.** Magnified view showing the catalytic triad of Subtilisin A bordering the DE loop of CVA9 VP1. S220 corresponds to the nucleophilic serine, D32 and H63 ensure the charge relay process of the catalytic site, and N154 stabilizes the attack through the oxyanion hole formation. For all predictions, the analyses were automatically stopped after 3 recycles. The experimental structures 1d4m (CVA9), 7c9y (CVB5), 1h8t (E11), 1scn (SubA, *B. licheniformis*) and 1s0q (BPT) have been used as template for all interface predictions.

**Figure 3:**
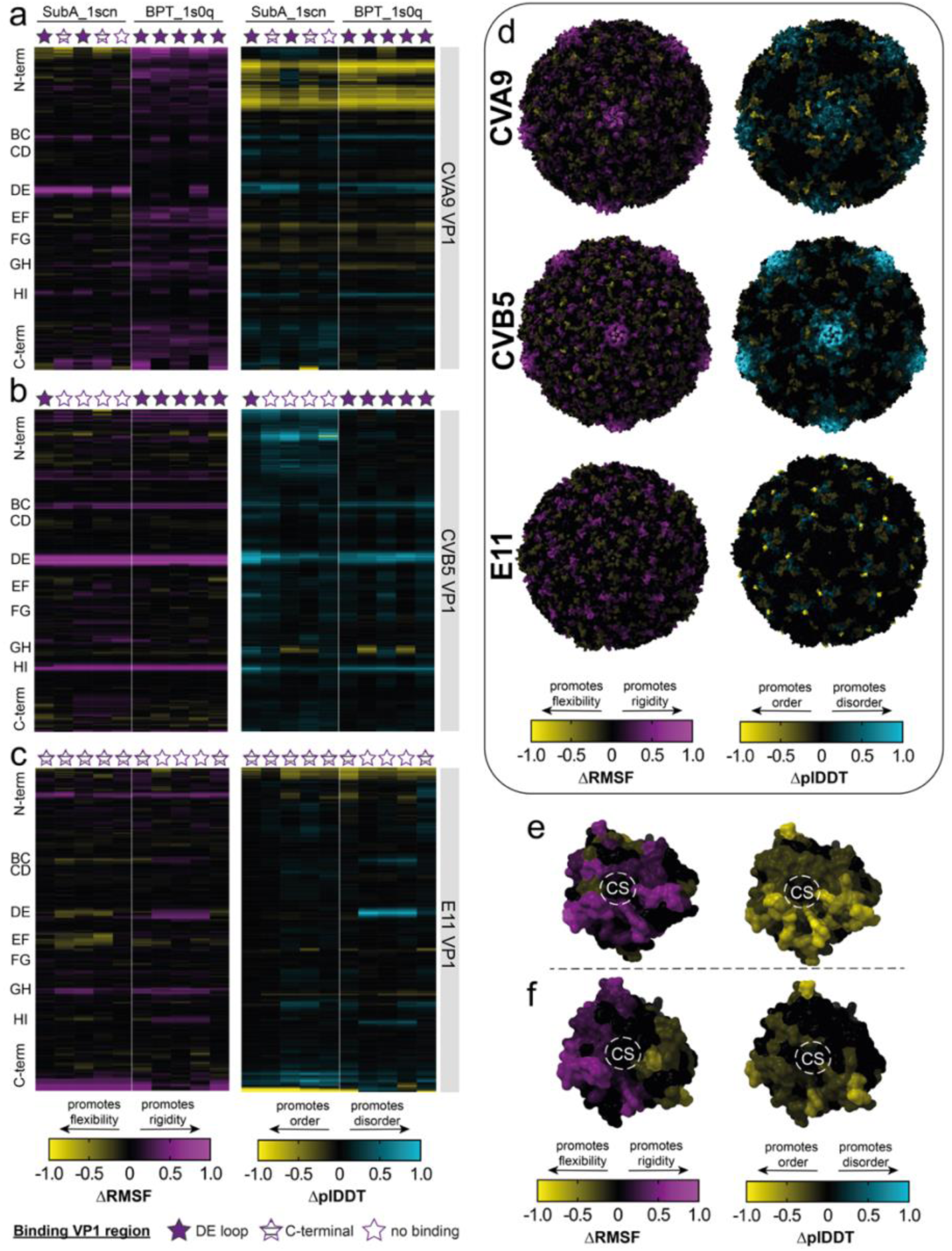
Both viral capsid IDPs and serine protease plasticity are challenged during protease/capsid exposures. **a-c.** Distribution of VP1 flexibility (*Δ*RMSF) or disorder (*Δ*plDDT) variations induced by Subtilisin A or BPT for **(a)** CVA9 **(b)** CVB5 and **(c)** E11. On the left of each heatmap, annotations correspond to the position of each IDPs in the viral sequence. On the top of each heatmap, stars indicate the predictive binding regions for each protease used, as mentioned above. On the top end of the first heatmap is indicated the name of the protease used in the analysis. **d.** Surface view of each viral capsid colored by *Δ*RMSF (left part) or by *Δ*plDDT (right part) for each virus type after an exposure with Subtilisin A. All the capsids have been centered on the 5-fold axis view. **e-f.** Surface view of Subtilisin A colored by *Δ*RMSF (left part) or by *Δ*plDDT (right part), after an exposure with **(e)** CVA9 protomer or **(f)** E11 protomer. (CS): catalytic site. All profiles variations have been calculated based on AF2-M models folded using 3 recycles.

**Figure 4:**
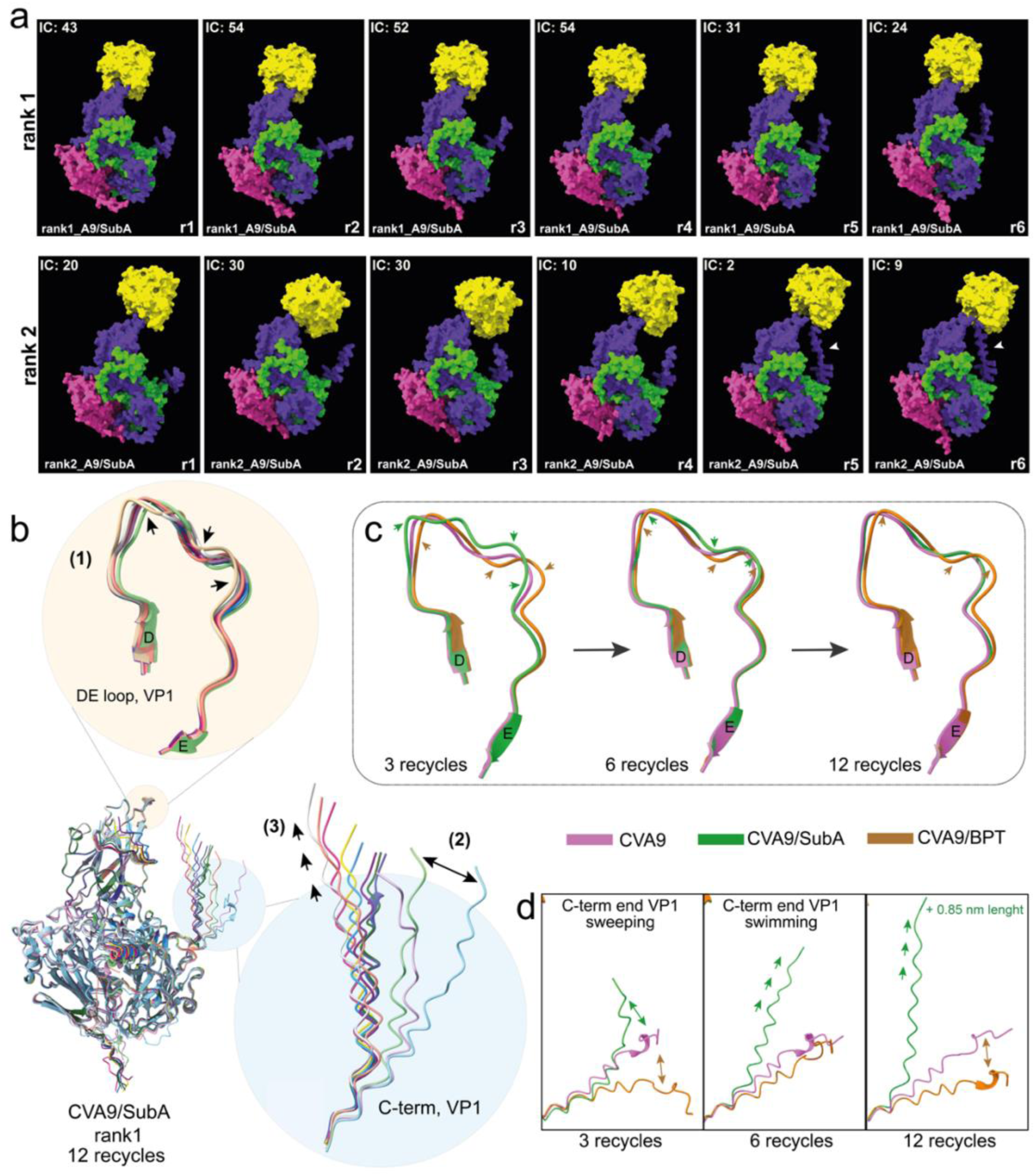
Subtilisin A induces swinging of CVA9 VP1 DE loop and leads to a fight-sensor swimming response of the C-terminal end VP1 sequence. **a.** Predictive timeline of Subtilisin A recruitment to CVA9 protomer. For the two ranks showing a binding to Subtilisin A, each timeline is composed of 6 recycles and the number of interatomic contacts (ICs) is specified in each frame. **b.** Analysis of the recruitment of Subtilisin A on CVA9 capsid by overlaying the CVA9 protomer modeled during 12 recycles in the presence of the enzyme. Subtilisin A was removed from the structural file to better assess predictive viral protein trajectories. On the upper part of the figure, (1) the swinging effect indicates the predictive trajectory induced by a binding with Subtilisin A to the VP1 DE loop. On the right part of the figure, (2) the sweeping effect and (3) the swimming effect correspond to the two sensor-response trajectories of the VP1 C-terminal end segment. **c.** Comparative trajectories induced by Subtilisin A and BPT binding on the VP1 DE loop of CVA9 after 3, 6, and 12 recycles. **d.** Comparative sensor-response trajectories observed after binding of the DE loop with Subtilisin A or BPT. On the right representation, 0.85 nm corresponds to the maximum additional distance swam by the VP1 C-terminal end segment (stretched form) after 12 recycles with Subtilisin A.

### Specific protease/virus molecular interactions

To determine the protein regions most likely involved in the interaction with serine proteases, we then used AF2-M to fold each viral protomer either with Subtilisin A or BPT. Since only Subtilisin A inactivates CVA9 in our biological experiment, we first investigated the recruitment of this protease on a CVA9 protomer. Among the top five predictive models returned by AF2-M, two showed an interaction with the DE loop of VP1, whereas two others targeted the C-terminal end VP1 sequence, all these IDPs being exposed on the surface of the capsid shell (Fig. 2 c-e, h, Extended data Fig. 5 a). While both CVB5 and E11 were insensitive to this protease, AF2-M nevertheless predicted an interaction between Subtilisin A and these virus types. One model returned for CVB5 (rank 1) indicated an interaction of this protease with the DE loop of VP1 (Fig. 2 f, h, Extended data Fig. 5 b), whereas an interaction with the C-terminal end of VP1 was predicted for E11 (rank 1 to 5) (Fig 2 g, h, Extended data Fig 5. c). We then reconducted this molecular modeling with each protomer and BPT, known to only cleave the sessile bond following an R found in tandem with the RGD motif of CVA9 C-terminal end^35^ (Extended data Fig. 3 c). The top five CVA9/BPT models indicated a protein contact between the protease and VP1 (Fig. 2 h). However, AF2-M did not predict the interaction of BPT with the C-terminal end sequence of VP1 but instead, it suggested the recruitment of this protease on the DE loop (Extended data Fig. 5 d). Comparable findings were obtained with CVB5 (Fig. 2 h, Extended data Fig. 5 e), while for E11 AF2-M predicted only the VP1 C-terminal end (embedding a K286) as the interacting region for BPT (Fig. 2 h, Extended data Fig. 5 f). The DE loop in all three virus types lacks a basic amino acid necessary for cleavage by BPT (Extended data Fig. 5 g-i), yet CVA9 and CVB5 allows an interaction with this protease. We therefore wanted to rule out the possibility of a protein attachment devoid of any predictable catalytic activity. However, for all models shown above, and regardless of the serine protease used for molecular modeling, the catalytic site was found in alignment with the viral substrate surface, supporting an attempt of enzymatic attack on those viral IDPs (Fig. 2 i).

**Figure 5:**
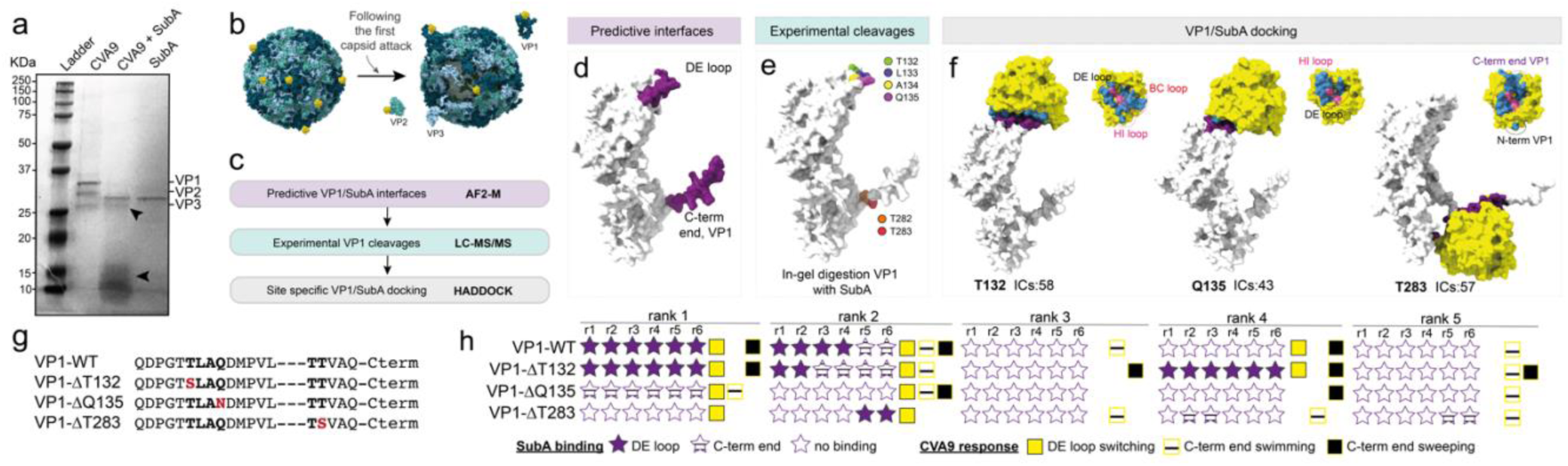
Three CVA9 VP1 amino acid residues contribute to the recruitment of Subtilisin A to CVA9 capsid and are structurally compatible to enzymatic cleavage. **a.** SDS-PAGE analysis of CVA9 capsid protein integrity following the digestion of purified native virus in solution with Subtilisin A. The two arrows point the proteolytic viral fragments observed after 15 min digestion with Subtilisin **b.** Schematic representation of the disintegration of CVA9 viral capsid following the predictive attack of VP1 on the 5-fold vertex and subsequent use of each independent viral protein as substrate. **c.** Deductive inference process used to identify amino acid residues involved in CVA9 inactivation by Subtilisin A. **d.** Surface representation of the two interfaces (in purple) predicted by AF2-M to interact with Subtilisin A during the early stages of CVA9 inactivation. **e.** Mass spectrometry identification of the six amino acid residues for which the C-terminal peptide bond was cleaved following in-gel digestion of denatured VP1 with Subtilisin A. **f.** Site-directed docking of Subtilisin A on three of six amino acid residues deduced during the two first steps of the selection process. Protein/peptide docking have been performed using HADDOCK 2.4 and interatomic contacts (IC) for each output have been determined using Prodigy. Surface views of Subtilisin A contact points with each VP1 regions are shown to the right of each docking structure. Pink surface regions correspond to the catalytic triad of the enzyme. **g.** Point mutations of VP1 applied *in-silico* and used to predict the significance of each amino acid residue in Subtilisin A recruitment on viral capsid. On the left of the dotted line, the sequence corresponds to the DE loop and on the right, the five first amino acid of the C-terminal end of VP1. **h.** Influence of CVA9 VP1 point mutations on the prediction of Subtilisin A recruitment to viral capsid. Subtilisin A recruitments on CVA9 protomer were predicted with AF2-M using 6 recycles. To the right of each prediction (rank 1 to 5), the squares represent the response of both DE loop and C-terminal end sequence trajectories. The absence of a square indicates that the corresponding trajectory event was not observed for any recycle of the analysis.

Since the recruitment of serine protease to viral IDPs was not explained by enzyme specificity, we next investigated if different predicted interactions established between the protein partners might be protease/virus specific. We therefore reconducted these experiments either by truncating the C-terminal end sequence of VP1s, by substituting the full DE loops with a hydrophobic VL repeat sequence or by challenging both proteases for the same protomer during analysis (Extended data Fig. 6 a-g). The presence of a hydrophobic DE loop in CVA9 (CVA9_DE loopΔVL) induced the partial or total loss of the interface between the proteases and the DE loop. However, each protease remained localized in the vicinity of this viral region during modeling, suggesting that the neighboring residues of this viral IDP also contribute to both proteases’ recruitment (Extended data Fig. 6 a-c). Conversely, for CVB5, we observed that replacing the DE loop with the same hydrophobic sequence favored the binding of Subtilisin A to this IDP, while reducing BPT binding to the same site (Extended data Fig. 6 e). For E11, the absence of a C-terminal end on VP1 does not help Subtilisin A to consider the DE loop as an interface, unlike BPT, for which the DE loop becomes the main target (Extended data Fig. 6 d). Finally, simultaneous exposure of each protomer with the two proteases revealed that BPT primarily and exclusively bound to viral DE loops, while Subtilisin A recognized the C-terminal end sequence of VP1 (Extended data Fig. 6 f-g). Consequently, the protease/capsid interactions modeled by AF2-M are both virus type and serine protease specific but are not predictors for viral inactivation.

### Long-range response of protease/capsid plasticity

IDPs are characterized by high levels of plasticity and constitute attractive targets for proteases because of their inability to adopt a stable three-dimensional structure^40,41^. While our first results indicated two specific viral IDPs as target of serine proteases, we wondered to what extent the computational presence of both protein partners predicts long-range structural fluctuations for all viral IDPs. As a first investigation of the early-stage recruitment of serine proteases on viral capsid, we monitored two structural indicators (Root Meaning Square Fluctuation (RMSF) and plDDT) after the computational exposure of each viral protomer to BPT and Subtilisin A. The degree of modification of both flexibility and disorder on each VP backbone, expressed by ΔRMSF and ΔplDDT respectively (Extended data fig. 7 a-b), led to identify distinct plasticity responses depending on the viral types and the proteases considered (Fig. 3 a-c, Extended data Fig.7 c-e). For VP1s, predicted to be the target of both serine proteases, the most apparent RMSF and plDDT fluctuations were found in three loops (BC, DE, HI) and in the N-/C-terminal regions of the protein (Fig. 3 a-c). However, the extent of such variation measured with both indicators appears to be greatest for CVB5 VP1 (Fig. 3 b), compared to CVA9 (Fig. 3 a) and E11 (Fig. 3 c), also suggesting that each viral type displays a specific plasticity fingerprint during computational exposure to each protease. In addition, no association between the prediction of a contact with the DE loop or the C-terminal end VP1 and the presence of fluctuations could be made, suggestive of long-distance interactions between both protein partners. Subtilisin A and BPT are both predicted to physically interact with the DE loop of CVA9, yet they do not induce the same structural fluctuation responses (ΔRMSF, ΔplDDT) on this IDP segment (Fig. 3 a). Conversely, for all E11 protomer models predicting a contact of Subtilisin A with the C-terminal end of VP1, more variable fluctuations in several loops were also found for both indicators (Fig. 3 c). Lastly, the analysis of these profiles for VP2 and VP3 from each viral type, which are not involved in the prediction of protease/capsid interfaces, also led to the identification of specific plasticity fingerprints for these two proteins (Extended data Fig. 7 c-e). To visualize these fluctuations on the entire capsids and thus better understand the protease/capsid structural variations, the ΔRMSF and ΔplDDT were projected onto the experimental assembly structures of each viral type (Fig. 3 d, Extended data Fig. 7 g). For CVA9 and CVB5 but not E11, Subtilisin A challenges the 5-fold vertex by concomitantly promoting a rigidification of the backbone and an increase in the local disorder of this region (VP1: BC, DE, HI loops) (Fig. 3 d). Comparatively, BPT induces a similar response around the 5-fold center of CVB5; however, the plasticity of the same region in E11 and CVA9 is considerably less challenged under similar conditions (Extended data Fig. 7 g). Finally, we applied the same methodology to study the structural fluctuations incurred by the two serine proteases when exposed to the three viral protomers (Fig. 3 e-f, Extended data Fig. 7 f, h, i). Similar ΔRMSF and ΔplDDT profiles were observed for each enzyme during exposure with each protomer, except for Subtilisin A during exposure with E11, which shows a less pronounced ΔplDDT profile than when exposed to CVA9 and CVB5 (Fig. 3 e-f, Extended data Fig. 7 f). Projecting these fluctuations onto the structures of each protease led to the identification of the region surrounding the catalytic sites of each enzyme as the most prone to structural shifts, though specific fingerprints could also be measured on each enzyme depending on the viral type exposed (Fig. 3 e-f, Extended data Fig. 7 h-i).

### Temporal evolution of protease/protomer binding

The conformational adaptation over time of a protease on a substrate is the prerequisite for any enzymatic cleavage. Consequently, we assumed that the temporal stability of protease/protomer complexes and the spatial mobility response of viral IDPs involved in these interactions might be discriminating predictors for viral inactivation. To investigate this concept, we used AF2-M as a timelapse simulator of the molecular recruitment of Subtilisin A and BPT on CVA9, CVB5 and E11 capsid protomers. Since adding iterative cycles throughout the network (‘recycles’) refines the structure of multimeric models^42^, we used this parameter to refine protease/virus interfaces. The number of recycles during the molecular modeling was increased (6 recycles) and we analyzed each recycling output as a unique time point of the prediction. During the recruitment of Subtilisin A on CVA9 protomer, only the first two ranks of the analysis predicted a contact between VP1 DE loop and this enzyme (Fig. 4 a). For the first timelapse (rank 1), the number of interatomic contacts (ICs) between Subtilisin A and the CVA9 vertex indicated a significant contact area during the first four recycles (IC_max_ = 54, r2), suggesting that in absence of effective cleavage, Subtilisin A continues to stably interact with this capsid region (Fig. 4 a, Extended data Fig. 8 a). In the second timelapse (rank 2), we also observed a contact between the DE loop and the enzyme, however the number of ICs was lower (IC_max_ = 30, r2), concomitantly leading to the recruitment of the enzyme by the C-terminal VP1 end segment and Subtilisin A unhooking from the DE loop (r5, IC=2) (Fig. 4 a, Extended data Fig. 8 b-c). We previously predicted the interface of Subtilisin A with the DE loop of CVB5 (Fig. 2 f) using a fast automated prediction set at 3 recycles, however this virus type was not inactivated in our biological assays (Fig. 1 b). By increasing the computational time (6 recycles) during the predictive recruitment of Subtilisin A to CVB5 protomer, we demonstrate that this protease interacts with the DE loop (rank 2), but this contact was likely too furtive (r1, IC_max_ =13) to correlate with any biological activity (Extended data Fig. 8 d). In contrast, a stable interaction with Subtilisin A was identified at the N-terminal end of VP2 for this virus type (rank 1), though this protein region is located inside the capsid and therefore not accessible in a full virus capsid. We finally confirmed that Subtilisin A does not interact with the DE loop of the 5-fold vertex of E11, but that the interaction with the C-terminal VP1 end of this viral type was maintained over time and could constitute a strong enzymatic trap for this protease (Extended data Fig. 8 e).

Though BPT did not induce inactivation for any of the three viruses, we previously predicted its interaction with the DE loop of CVA9 and CVB5 (Fig 2. h). To confirm the robustness of these predictions, the 6-recycles modeling protocol was applied for all three virus types with BPT and confirm the initial predictions (Extended data Fig. 8 f-h). More specifically, the interaction of BPT with VP1 DE loop in CVA9 describes the prediction of 32 ICs after 4 recycles, this interaction being maintained during the last three recycles of the analysis. For CVB5, we noted a greater number of ICs with the VP1 DE loop (IC_max_=59, r2) than for CVA9 VP1 DE loop (IC_max_=32, r4), suggesting that BPT binds more strongly to the vertex of CVB5, yet is not able to use this protein portion as a substrate. Finally, for E11, the molecular modeling confirmed the interaction between BPT and the C-terminal end of VP1. Therefore, all the results obtained in the presence of BPT strongly predict a trapping of the enzyme on the three viral types, which does not lead to virus inactivation. While increasing recycling strengthens predictions of potential interfaces and eliminates weaker ones, it remains however imprecise to predict virus inactivation due to the trapping of some proteases on non-functional viral IDPs.

### CVA9 sensor response induced by Subtilisin A

Since increasing the number of recycles strengthened the previous virus/protease interface predictions, we finally used this strategy to refine the understanding of the molecular recruitment of Subtilisin A on CVA9, which leads to virus inactivation. Besides the interaction with the DE loop, analyses conducted with 6 recycles indicated the recruitment of Subtilisin A by the C-terminal end VP1 sequence, this phenotype not being observed with BPT. Therefore, we hypothesized that the local recruitment of the C-terminal end sequence to the DE loop binding Subtilisin A might be a defensive response to the enzymatic cleavage hazard pressure on this IDP. Molecular timelapse prediction of Subtilisin A recruitment to CVA9 protomer were thus reconducted by increasing the computational time up to 12 recycles. The resulting captures indicate that the C-terminal end VP1 sequence is predicted to act both as a DE loop competitor substrate (SI Video 1) or as a DE loop protective peptide (SI Video 2), both mechanisms ultimately leading to 5-fold vertex protection. To better visualize the spatial mobility of viral protein elements during predictive enzymatic attack, the structures resulting from each recycle were overlaid (rank 1, n_recycles_ =12) (Fig 4. b). While the DE loop shows a swinging motion (1) in response to a physical contact with Subtilisin A, the C-terminal end VP1 sequence, which shows a sweeping trajectory during the first recycles (2), seems to be quickly forced to adopt a swimming motion towards the 5-fold vertex (3). To illustrate that this virus response was exclusively encountered in the presence of Subtilisin A, we overlaid the structures of CVA9 modeled with or without BPT or Subtilisin A to compare the trajectories of these regions of interest over 12 recycles (Fig. 5 c, d). After the first recycle, the mobilization of the DE loop by BPT and Subtilisin A results in a strong misalignment of this viral IDP, as compared to the position taken by the loop backbone in absence of protease (Fig. 5 c). After 6 recycles, a better alignment of the DE loop with both proteases was observed, compared to the condition without protease. Finally, after 12 recycles, no difference in DE loop trajectory was observed between the modeling without protease and the one with Subtilisin A, indicating that the recruitment of the enzyme by the C-terminal end sequence of VP1 allows the DE loop to recover its native position. For BPT, no visual difference was observed following 12 recycles of analysis compared to the models obtained after 6 recycles, suggesting that the presence of the BPT freezes the spatial mobility of the DE loop over time, and this regardless of the presence of cleavage site on the loop. Similarly, we analyzed the different trajectories taken by the C-terminal end VP1 sequence in presence of serine proteases in the computational environment. During the initial recycles of the analysis mimicking the contact of both Subtilisin A and BPT with the VP1 DE loop, we observed that this IDP adopted a sweeping motion, this trajectory being not observed in absence of protease (Fig. 4 d). Whereas this trajectory was maintained in the presence of BPT during higher recycles, the C-terminal end sequence took a completely different path following the contact of Subtilisin A with VP1 DE loop. Indeed, after 6 recycles, the C-terminal end segment began a progressive swim towards the center of the 5-fold vertex of the virus capsid, this process resulting in an additional elongation of the C-terminal end sequence (up to 0.85 nm), sufficient to unhook the protease from the DE loop (Fig 4. D, SI video 2). Therefore, these predictions suggest that the enzymatic pressure of Subtilisin A on the DE loop most likely acts as a sensor to drive the protective recruitment of the C-terminal end VP1 sequence to the 5-fold vertex of CVA9 capsid.

### Site-specific inactivation of CVA9 by Subtilisin A

To resolve the identification of cleavages on capsid regions leading to viral inactivation, we finally compared experimentally determined cleavage sites with predictive structural ones obtained for CVA9 following an exposure to Subtilisin A. We first treated purified CVA9 virions with Subtilisin A before analyzing all capsid degradation products by SDS-PAGE (Fig. 5 a). The treatment induces a distinctive loss of the two bands corresponding to proteins VP1 (33.8 KDa) and VP2 (28.4 KDa) (CVA9 + SubA), providing evidence for the inactivation of CVA9 by capsid degradation and indicating that both VP1 and VP2 are ultimately involved in the enzymatic race dynamics of Subtilisin A (Fig. 5 a,b). Findings were less conclusive for the VP3 protein, where exposure to Subtilisin A yielded a band at 26.5 KDa, very close to the expected mass of VP3 (26.3 KDa) (Fig. 5 a). Since VP1 was confirmed to be cleaved by Subtilisin A treatment, we subsequently used a deductive inference process to trace the effective cleavage sites of the enzyme on this protein (Fig. 5 c). We used the two interfaces predicted by AF2-M as a coarse screening of Subtilisin A cleavage sites (Fig. 5 d) before confirming these predictions by protein mass spectrometry analysis. Finally, to assess enzyme/substrate accessibility during viral inactivation, each experimentally confirmed cleavage site was subjected to a site-specific molecular docking analysis with Subtilisin A. Of the 80 cleavage sites identified after in-gel digestion of VP1 with Subtilisin A, four were identified on the DE loop (T132, L133, A134, Q135) and two on the C-terminal end part of the protein (T282, T283) (Fig. 5 e, Extended data Fig. 9 a-d.). To discriminate whether cleavage of this VP in denatured form could also occur in a folded form, we used VP1 and Subtilisin A protein structures modeled by AF2-M to generate site-directed molecular docking between each residue preceding a cleavage event by Subtilisin A and the catalytic site residues (H32, D63, N154, S220) (Extended data Fig. 9 e). For the six residues identified as potential targets of Subtilisin A on CVA9, three of them (T132, Q135, T283), showed a distance of less than 3 Å with the nucleophilic serine of Subtilisin A (S220), indicative of hydrogen bonding between the two partners (Fig. 5 f, Extended data Fig. 9 f). For each refined docking, we also analyzed the contact areas between the enzyme and the viral substrate, which indicated a number of ICs ranging from 43 to 57 depending on the sites of action (Fig. 5 f). Previously, we found that the BC and HI loops may contribute to the recruitment of Subtilisin A to CVA9 (Fig. 3, Extended data Fig. 8 a). We indeed noticed that these two loops also physically interact with the enzyme during docking (Fig. 5 f, Extended data Fig. 10 a). Specifically, the docking targeting T132 indicates that Subtilisin A uses the HI and BC loops of VP1 to stabilize the protease/VP1 complex. For docking targeting Q135, the HI loop of VP1 also provides a support for the enzyme. Moreover, the overlay of Subtilisin A-binding VP1 on a CVA9 VP1-pentamer indicates that loops of adjacent VP1s also contribute to protease fitting on the 5-fold vertex (Extended data Fig. 10 b). Finally, for the docking of Subtilisin A on the C-terminal end sequence (T283), we identified the N-terminal region of VP1 and a part of VP3 surface as additional contact area for interaction (Fig. 5 f, Extended data Fig. 10 c). During the attack on the three sites most likely responsible for CVA9 capsid degradation, Subtilisin A therefore fits other capsid regions before proceeding with enzymatic cleavage.

### Site-specific viral response

To finally gain an understanding of the role of each residue targeted by Subtilisin A in the viral response, we generated *in-silico* substitutions on each residue of interest before predicting the phenotypic effect of each mutation with AF2-M. To preserve the hydropathy of the viral regions as much as possible, each amino acid residue was replaced by a residue with a similar side chain and with the closest hydropathy index (H_i_) of the original residue (Fig 5 g, Extended data Fig. 10 d). The predictive impact of mutations T132ΔS132 and Q135ΔN135 led to the identification of two different responses in presence of Subtilisin A (Fig. 5 h). Specifically, we noticed, in comparison with VP1-WT, that a substitution of T132 (VP1-ΔT132) did not cause any measurable negative effect on the ability to predict the DE loop and the C-terminal end sequence as interfaces for Subtilisin A, and similar conclusions were obtained regarding the response trajectories of both protein moieties. Conversely, we observed that the Q135ΔN135 (VP1ΔQ135) mutation led to the complete loss of prediction of the DE loop as an interface for the enzyme, thus suggesting that this residue plays a crucial role in enzyme/substrate recognition. However, this mutation does not prevent Subtilisin A from considering the C-terminal end VP1 sequence as an interface, nor does it appear to abrogate the different types of viral mobility responses (Fig. 5 h). Similar analyses were conducted with residues L133 (L133ΔI133) and A134 (A134ΔV134), which are experimentally targeted by Subtilisin A using denatured VP1 but are not structurally accessible on the native form of the protein (Extended data Fig. 10 d-f). The predictive results returned for these two conditions maintain the DE loop and the C-terminal end sequence as interfaces for Subtilisin A and demonstrate similar mobility responses, thus contributing to discredit these two residues in the early stages of enzyme/substrate recognition. Finally, a similar approach was conducted on residues T282 and T283 (C-terminal end VP1), for which the peptide bond is cleaved by Subtilisin A, though only T283 indicated to be structurally compatible for cleavage (Fig. 5 g, Extended data Fig. 10 d). For the VP1-T283 mutant (T283ΔS283), we observed a partial loss of recognition of the DE loop as interface for Subtilisin A, compared to the VP1-WT condition. However, no change was observed regarding the recognition of the C-terminal end of VP1 by this protease (Fig. 5 h). We also noticed that the mobility of the DE loop was conserved, as well as the ability of the C-terminal end sequence to adapt an extended shape (swimming trajectory). However, the mutation T283ΔS283 no longer allows the sweeping motion of the C-terminal end sequence, suggesting that this trajectory contribute very likely to the DE loop mobility during the Subtilisin A recognition process. For VP1-T282 mutant (T282ΔS282), we observed some variability in the prediction of the C-terminal end sequence as interface, but this mutation doesn’t abrogate the ability of Subtilisin A to consider the DE loop as interface (Extended data Fig. 10 e). Furthermore, no change in the mobility response of VP1-IDPs was observed with VP1-T282, indicating that this residue is poorly involved in DE loop swinging effect, as well as in multi-axial movements of the VP1 C-terminal end sequence. Therefore, of the two CVA9 DE loop residues targeted by Subtilisin A that most-likely lead to viral inactivation, one of them (Q135) is also crucial in the protease’s recognition process on the 5-fold vertex. Furthermore, T283, found on the C-terminal end VP1 sequence, is the initiator residue of the sweeping motion, thereby acting as a sensor residue to facilitate enzyme recognition on the DE loop.

## Discussion

While the repertoire of extracellular bacterial serine proteases found in aqueous environments contributes to the extra-host inactivation of enteroviruses^3^, the underlying molecular mechanisms explaining such an outcome remained unexplored. Accordingly, we provided here a first overview of the structure/function relationship between these enzymes and miniatures versions of enterovirus capsids (protomer; capsid portion: 1/60^th^), thereby leading to a better understanding of protease-mediated inactivation.

For the three virus types used in this work (CVA9, CVB5, E11), only CVA9 was sensitive to both serine proteases studied (Subtilisin A, BPT), corroborating with previous data which suggest an atypical sensitivity of this virus type to proteolytic enzymes^4^. Furthermore, we demonstrated that both proteases cleave the C-terminal end sequence of CVA9 without altering virus infectivity^35,36^, but only Subtilisin A inactivate CVA9 in a host-independent manner. These experimental findings can be rationalized by an integrative computational approach, combining the prediction of protease/capsid interfaces and a site-specific docking refinement. Consequently, the absence of a stable interface between Subtilisin A and the DE loops of the 5-fold vertex of CVB5 and E11 explains the proteolytic resistance of both viruses. Conversely, CVA9 DE loops predict an interface with Subtilisin A, and the molecular docking refinement targeting two loop residues explains CVA9 sensitivity to inactivation in a host-independent manner. Predictions of interfaces with BPT for all viruses indicate that DE loops interact with this enzyme, yet the three viruses are insensitive to proteolysis in a host-independent manner due to the absence of a specific cleavage site (K, R). Finally, site-directed docking analyses of CVA9 C-terminal end residues (T283: Subtilisin A; R288: BPT (data not shown)) explain the cleavage by both proteases, leading to the loss of the RGD motif used for integrin recognition in BGMK cells.

While viral inactivation mimicked the reactional endpoint of enzyme catalysis, the intermediate steps leading to such a phenotype remained to be demonstrated. Thanks to the release of AF2-M^38^, we reconstructed with accuracy enterovirus capsid protomers, subsequently facilitating the functional study of protease/capsid complexes by molecular modeling. By selecting CVA9 as a serine protease sensitive virus and CVB5 and E11 as insensitive viruses to challenge our hypotheses, we sequentially reassembled the molecular steps leading to enzymatic cleavage of viral capsids. As such, we propose a three-step predictive molecular mechanism to explain the recruitment of these enzymes on such giant substrates:

**(1) Precognition** - Before any physical contact between serine proteases and viruses, the co-presence of both partners in the same environment led to structural fluctuations highly predictive of long-range electrostatic interactions^43^. These interactions have already been described as a modulator of folding stability of IDPs^44,45^, and contribute to the determinism and stability of serine proteases for substrates^46,47^ as well as to the functional modulation of some allosteric enzymes^48,49^. For each modeling performed with serine proteases and viral protomers, we mainly observed those fluctuations on viral IDPs and on regions surrounding the active sites of each protease, which is highly consistent with the findings cited above. Moreover, we point out different flexibility responses depending on the partners used, also suggesting that long-range interactions are both protease and viral substrate specific. Therefore, we suggest the contribution of long-range electrostatic interactions as an early stage of enzyme-substrate communication, which modulate both viral capsid and protease plasticity.
**(2) Recognition** - As a second step of interaction, we described the physical contact of serine proteases to some accessible IDPs segments of viral capsids (VP1: DE loop, C-terminal end). Due to the inability of IDPs to adopt a well define structural conformation, those sequences are known to depict disorder-to-order transitions states upon binding^50^, making them also sensitive to proteolysis^51^. Though viral IDPs have not been described as potential target of serine proteases, two studies indicate that both subtilisins and trypsins may favor the cleavage of accessible disorder segments in other native proteins^41,52^. However, we noticed that AF2-M prediction of protease binding to viral IDPs was not a binary indicator of enzymatic cleavage, but more an essential first contact to initiate a functional or a non-functional enzyme-substrate binding. Indeed, two facts pointing in this direction have emerged from this work, including the long-term binding of BPT to both CVA9 and CVB5 VP1 DE loops containing no effective cleavage site, and the brief binding of Subtilisin A to the DE loop of the CVB5 capsid, neither of which resulted in protein cleavage. While increasing the number of recycles during modeling can be valuable to refine structure prediction^53^, the modulation of this parameter also brings robustness to temporality and strength of protease/capsid interfaces. As a result, the interface of Subtilisin A with CVB5 DE loop was abrogated, but the non-functional binding of BPT with the DE loop of capsids was maintained. We therefore propose that this recognition step does not depend on the specificity of action of serine proteases for viral substrates, but rather on the strength of short-range interactions (*i.e.* van der Waals forces, hydrogen bonding) between the two protein partners, which further accommodates the ‘fight or flight’ response of serine proteases on viral capsids.
**(3) Adaptation** - Finally, the last step of enzyme/substrate communication achieved by site-specific molecular docking refinement mimics a protease/capsid structural adaptation, which is decisive for initiating any enzymatic cleavage. Though the active site of Subtilisin A physically binds the two CVA9 VP1 IDPs segments cleaved by the protease (DE loop, C-terminal end), the enzyme also adapts locally to other regions of the viral capsid to initiate catalysis. Indeed, the C-terminal end portion of CVA9 VP1, which seems to act as a ‘crazy horse’ with a great ability to elongate and move in space, fits locally on either side of the active site on the 5/2 axis of the capsid shell during molecular docking refinement. Similarly, the DE loop targeted by Subtilisin A, adapts its positioning on either side of the catalytic center with the adjacent BC and HI loops, these loops being described to contribute greatly to the plasticity of the 5-fold vertex of enteroviruses capsids^10,54^. Accordingly, we suggest that the adaptation step, which corresponds to the discriminative phase of protease-capsid interaction, clearly approximates the induced-fit model proposed by D. Koshland^55^.

While the presence of single mutations in a protein sequence is described to have minimal impact on the folding predicted by AlphaFold and on the overall stability of proteins^56^, our results indicate however that such mutations can modulate both protease/capsid interfaces prediction. For all single mutations generated *in silico* on the three effective cleavage sites of Subtilisin A on CVA9 VP1, AF2-M predicted a partial or complete loss of contact of the enzyme with the capsid VP1 DE loop depending on the viral mutant. Moreover, we also observed spatial mobility alterations of the C-terminal end VP1 sequence, thus suggesting that the effective residues of the viral substrate are involved in both the early recruitment of the enzyme on the capsid and the VP1 structural mobility response. Consequently, though the general recognition process of serine proteases on enterovirus capsids seems broadly similar, we point out that the side-chain chemistry of the targeted residues during catalysis, and by extension the predictable intermolecular forces engaged by these residues in the interaction with Subtilisin A, contribute to the specific interaction protease/capsid leading to viral inactivation. Overall, we provide a first methodological framework to investigate the functional characterization of serine proteases involved in the race for the disintegration of non-enveloped viruses in aqueous environments. Due to the high number of enzymatic reactions described to date and their roles in systems biology (*e.g.* microbial control, cancer research, food preservation), this advance will more generally speed up the knowledge on molecular mechanisms of protease/substrate interactions.

## Methods

### Cells and viruses

Buffalo Green Monkey Kidney (BGMK, provided by Spiez Laboratory, Switzerland) cells were maintained in minimum essential media (MEM, Gibco, ThermoFisher Scientific) supplemented with 10% Fetal Bovine Serum (FBS, Gibco) and 1% Penicillin/Streptomycin (P/S, Gibco). The human Rhabdomyosarcoma muscle cell line (RD, ATCC CCL-136) was maintained in Dulbecco’s Modified Eagles Medium (DMEM, Gibco) containing 10% FBS and 1% P/S. Both cell lines were maintained at 37°C (5% CO_2_). The environmental isolate coxsackievirus A9 (CVA9), originating from sewage, was provided by the Finnish National Institute for Health and Welfare. Echovirus 11 (E11) and coxsackievirus B5 (CVB5) were purchased from ATCC and correspond to the Gregory strain (VR-737) and the Faulkner strain (VR-185), respectively.

### Virus stocks preparation and enumeration

Each virus type was propagated by infecting sub-confluent monolayers of BGMK cells as previously described^1^. Briefly, viruses were released from infected cells following three freezing-thawing steps. Cell debris were removed by centrifugation (3’000 *x g*, 5 minutes). After filtration of the supernatant containing viruses on a 0.2 μm syringe filter (Filtropur S, PES, Sarstedt), virus stocks were concentrated on a 100 KDa cellulose membrane (Amicon Ultra-15, Merck Millipore) and rinsed three times successively with PBS. Infectious virus concentrations were enumerated by a most probable number (MPN) infectivity assay. Virus sample aliquots (100 μL) were diluted by 10^-1^ to 10^-8^ in MEM 2% FBS. Each dilution (100 μL, 5 replicates) was plated on a BGMK sub-confluent monolayer in a 96-well plate and was incubated at 37°C (5% CO_2_). After 4 days, the number of wells with cytopathic effects (CPEs) for each dilution was recorded, and the resulting number of infectious viruses per sample was calculated using R^2^. The limit of detection (LoD) of the assay, defined as the concentration corresponding to one positive cytopathic effect in the lowest dilution of the MPN assay under the experimental conditions used, corresponds to 2 MPN/mL. Each virus stock was stored at −20°C until use.

### Protease-mediated viral infectivity reduction assay

To screen the sensitivity of CVB5, CVA9 and E11 to serine proteases, 100 μL of each virus stock (1.10^6^ MPN/mL) were incubated with a final concentration of 20 μg/mL of Subtilisin A (P5380, Sigma-Aldrich) or Bovine Pancreatic Trypsin (BPT) (T1426, Sigma-Aldrich) for 2 or 6 hours at 37°C. All enzymatic reactions were immediately stopped by adding 900 μL of MEM 2% FBS. Negative controls of inactivation for each virus type were performed by replacing serine proteases by PBS, following the same procedure as described above. Viral reduction infectivity was determined as log_10_ (C/C_0_), where C is the residual titer after adding protease for each time point, and C_0_, the titer measured without addition of protease for each corresponding time point. The experimental LoD of the assay was approximately 5-log_10_.

### Flow cytometry analysis

Flow cytometry was used to determined αVβ3 and αVβ6 integrins surface expression in BGMK and RD cells. After a wash with PBS, cells were detached using trypsin-EDTA (0.05%) (Gibco) and pelleted by centrifugation (400 *x g*, 2 min) in a 96-well U-bottom plate (10^5^ cells/well). Cells were washed twice in PBS and stained with 1 μg/mL DAPI (D9542, Sigma-Aldrich) for 15 min at room temperature (RT) in the dark. After two successive washes with a staining buffer (PBS, 1% bovine serum albumin), cells were incubated with a Fc receptor blocking solution (564219, BD Pharmingen) for 15 min at RT in the dark. Subsequently, cells were stained with Alexa Fluor® 488-conjugated anti-αVβ3 antibody (Novus Biologicals, USA) (39.5 μg/mL) and Alexa Fluor® 700-conjugated anti-αVβ6 antibody (Novus Biologicals) (30.5 μg/mL), or with isotype controls (Alexa Fluor® 488-conjugated mouse IgG1 (R&D systems, USA) and Alexa Fluor® 700-conjugated rat IgG2 (Novus Biologicals) for 20 min at 4℃ in the dark. Stained cells were finally washed twice with PBS and resuspended with 200 μL PBS prior to an immediate acquisition. Data acquisitions were performed on a Gallios flow cytometer (Beckman Coulter, CA, USA), with a minimum of 5000 cells acquired per sample. Acquired data were analyzed in FloJo software version 10.8.0. Cell doublets were excluded by single cell gating, and single cells were then gated based on viability (DAPI^-^) prior to the analysis of cell surface integrin expression (αVβ3^+^ / αVβ6^+^).

### CVA9 C-terminal end VP1 cleavage assessment by cell culture

BGMK and RD cells were used to monitor the cleavage of CVA9 C-terminal end segment carrying a RGD motif. First, to investigate the permissiveness of both cell lines for CVA9, roughly 10^5^ MPN/mL of viruses were used to infect simultaneously 96-well plates of sub-confluent RD and BGMK monolayers following the general procedure as described above. The CPEs were monitored 3, 4 and 5 days-post infection and virus titers were calculated as mentioned above. To assess the cleavage of the C-terminal end VP1 sequence by Subtilisin A, two tubes each filled with 100 μL of CVA9 stock (1.10^6^ MPN/mL) were incubated with 2 μL of Subtilisin A at 1 mg/mL for 2 hours at 37°C. After incubation, all tubes were immediately filled with 900 μL of MEM 2% to stop the enzymatic reaction. The same incubation was similarly conducted with BPT and negative controls of inactivation were performed by replacing serine proteases by PBS. All conditions were simultaneously plated either on RD or BGMK sub-confluent monolayers as mentioned above. CPEs were monitored 5-days post infection and viral decays were calculated as described above. The experimental LoD of the assay was approximately 5-log_10_.

### Virus purification

Three T-150 cell culture flasks (25 mL/flask) of CVA9-infected BGMK cells were recovered and subjected to three successive freeze/thaw cycles. The cultures were centrifuged at 3’000 *x g* for 8 minutes and the resulting supernatants were filtered through 0.22 µm membrane. A volume of 30 mL of supernatant was dispensed in two 38.5 mL ultracentrifuge tubes (Ultra-Clear, 25×89 mm, 344058, Beckman Coulter) before adding 5 mL of a 20% sucrose solution in NTE buffer (100 mM NaCl, 10 mM Tris-HCl pH7.5, 1 mM EDTA pH8,) to the bottom of each tube. Each tube was centrifuged for 3 hours at 150’000 *x g* (4°C) using an Optima XPN-80 ultracentrifuge (Beckman Coulter) equipped with a SW 32 Ti swinging bucket (369650, Beckman Coulter). Supernatants were discarded by inverting tubes, and 80 µL of PBS was added to each individual pellet. After resuspension of pellets, the two samples were mixed, and the total volume of viral suspension was adjusted to 1 mL with PBS.

After packing a seven-phases discontinuous sucrose gradient (15%, 20%, 25%, 30%, 35%, 40%, 50%) in a 17 mL ultracentrifuge tube (Ultra-Clear, 16×102 mm, 344061, Beckman Coulter), and leaving overnight to allow for merging of the layers into a continuous gradient, 500 µL of CVA9 suspension were applied on top of the first layer. The tube was centrifuged for 3 hours at 150’000 *x g* (4°C) using a SW 32.1 Ti swinging bucket (369651, Beckman Coulter). Finally, successive 500 µL fractions were collected from the bottom of the tube and stored at −20°C until use.

### SDS-PAGE analyses

SDS-PAGE was used to monitor both CVA9 purity following the two-step sucrose purification procedure and to evaluate the proteolytic degradation of viral proteins (VPs) by Subtilisin A. Prior to each SDS-PAGE analysis, 20 µL of protein sample were denatured in Laemlii buffer for 10 min at 95°C. The protein profiles in each sucrose fraction and the identification of the purified CVA9 fraction have been analyzed on Any KD Mini-Protean TGX Precast gel (4569033, Biorad). Sample preparation of CVA9 VPs for in-gel digestion prior to mass spectrometry analyses and the assessment of the VPs degradation following Subtilisin A treatment on native CVA9 virions have been done using 12% Mini-Protean TGX precast gel (4568043, Biorad). Prior to SDS-PAGE analysis of VPs degradation products by Subtilisin A, 20 µL of the purified native CVA9 fraction were incubated either with 2 µL of Subtilisin A at 100 μg/mL or with 2 µL of PBS for 15 minutes at RT. For all analyses, the Dual Color standards (Biorad) has been used as protein ladder and gels were stained with Instant Blue Coomassie solution (Abcam).

### In-Gel digestions

Gel pieces containing the concentrated individual VPs were excised and washed twice with 50% ethanol in 50 mM ammonium bicarbonate (AB, Sigma-Aldrich) for 20 min and dried by vacuum centrifugation. Proteins were reduced with 10 mM dithioerythritol (Merck-Millipore) for 1 h at 56°C followed by a washing-drying step as described above. Reduced proteins were alkylated with 55 mM Iodoacetamide (Sigma-Aldrich) for 45 min at 37°C in the dark followed by the same washing-drying step described above. VPs were then digested either (1) overnight at 37°C using mass spectrometry grade Trypsin (Pierce), GluC (Pierce) or Chymotrypsin (Promega) at a concentration of 12.5 ng/µl in 50 mM AB, or (2) for 1 hour at 37°C using Subtilisin A (P5380, Sigma-Aldrich) with the same concentration as for the three other enzymes. For Trypsin digestions, 10 mM CaCl2 was added. Resulting peptides were extracted in 70% ethanol, 5% formic acid (FA, Merck-Millipore) twice for 20 min, dried by vacuum centrifugation and finally desalted on C18 StageTips^3^.

### Mass spectrometry analyses

Digested-VPs were resuspended in 2% acetonitrile (ACN, Biosolve), 0.1% FA for LC-MS/MS injections. Nano-flow separations were performed on a Dionex Ultimate 3000 RSLC nano UPLC system (Thermo Fischer Scientific) online connected with an Orbitrap Lumos Fusion Mass-Spectrometer (Thermo Fischer Scientific). A capillary precolumn (Acclaim Pepmap C18, 3 μm-100Å, 2 cm x 75 μm ID) was used for sample trapping and cleaning. Analytical separations were conducted on a 50 cm long in-house packed capillary column (75 μm ID, ReproSil-Pur C18-AQ 1.9 μm silica beads, Dr. Maisch). The analysis was performed with a 250 nL/min flow rate (A: 98% H_2_O, 2% ACN, 0.1% FA; B: 90% ACN, 2% H_2_O, 0.1% FA) using a 90 min biphasic gradient as follow: after sample loading at 1% B, the gradient was raised to 24% B in 46 min, followed by an increase to 38% B in 10 min. After 3 min at 90% B, a final conditioning was set at 1% B for 15 min. Acquisitions were performed through Top Speed Data-Dependent acquisition mode using a cycle time of 1 second. First MS scans were acquired over a mass range of 375 to 1’500 m/z with a resolution of 240’000 (at 200 m/z) and a maximum injection time of 50 milliseconds was used. The most intense parent ions were selected and fragmented by High energy Collision Dissociation (HCD) with a Normalized Collision Energy (NCE) of 30% using an isolation window of 0.7 m/z. Fragmented ions were acquired using the Ion Trap with a maximum injection time of 60 milliseconds. Selected ions were then excluded for the following 20 seconds. Raw data were processed using SEQUEST and MS Amanda^4^ in Proteome Discoverer v.2.2 against the Uniprot Chlorocebus aethiops entries supplemented with VP1, VP2 and VP3 ones (from PDB 1d4m). Enzyme specificity was set either to Trypsin, GluC or Chymotrypsin and a minimum of six amino acids was required for peptide identification. Up to two missed cleavages were allowed and a 1% FDR cut-off was applied both at peptide and protein identification levels. Data were further processed and inspected in Scaffold 5 (Proteome Software, Portland, USA).

### Multimeric protein folding

Both viral protomers folding and protease/virus interfaces were predicted with AlphaFold2-Multimer v2 (AF2-M)^5^ using ColabFold v1.5.2^6^. Multiple sequence alignments (MSA) were generated with MMseqs2^7^ against the Uniclust30 database^8^, the MGnify database^9^ and the PBD70 database^10^ using the unpaired_paired mode. For each protein sequence inputs used for analysis, PDB templates were added to prediction: CVA9 (1d4m), E11 (1h8t), CVB5 (7c9y), Subtilisin A (1scn), BPT (1s0q). The folding of each viral protomer and the fast prediction of protease/capsid interfaces were initially performed in automatic mode using 3 recycles. To challenge the strength of protease/capsid interfaces and study the long-term binding of proteins partners, the number of recycle has been increased up to 6 or 12 depending on the experiment with an early step tolerance set at 0. Topologies qualities of the folded proteins and interfaces were estimated using Template-Modeling (TM) score, interface-TM score, predicted local Distance Difference Test (plDDT) and Predicted Alignment Error (PAE) plot. The model confidence of each folded monomer or multimer was calculated as previously described^5,11^. A comparison of the overall RMSD between predictive viral protomers and the corresponding PDB structure was also used for the analysis. For the timelapse simulator experiment of protomer/protease interfaces (n_recycles_ = 6), the PRODIGY webserver^12^ was used to assess the number of Interatomic Contacts (ICs) between each protein partner for each recycle.

### Plasticity response assessment

To assess the local plasticity response of both viral protomers and proteases during the folding process, pLDDT and Root Meaning Square Fluctuation (RMSF) values were used as structural motion indicators. Both indicators were studied on all predictive protease/protomer models and compared with a predictive protease model alone or a predictive protomer alone depending on the interaction model. As such, disorder assessment was expressed as ΔplDDT = plDDT_0_ – plDDT_1_, whereby plDDT_0_ corresponds to the global order/disorder state of either a predicted protease or a predicted protomer without encountering the other protein partner and plDDT_1_ defined the plDDT output of the co-presence of both protomer and protease in the computational environment. To assess flexibility variations, the CABS-flex 2.0 standalone webserver^13^ was run with the same PDB files returned by AF2-M as used above. Flexibility variations were calculated as ΔRMSF = RMSF_0_ – RMSF_1_, whereby RMSF_0_ corresponds to the global flexibility/rigidity state of each protein partner alone and RMSF_1_ corresponds to the same profile during the computational exposure of both protein partners. All Δvalues, initially ranging from −40 to 40 for ΔplDDT and −8 to 8 Å for ΔRMSF depending on protomer/protease models, have been normalized between -1 to 1, whereby 0 indicate no detectable plasticity change during the folding process.

### Predictive viral mutants

To assess interface competition between each protease with the DE loop and the C-terminal end of VP1 during the folding with E11 and CVA9, the VP1 sequence of each viral type was modified either by truncation or substitution. Truncated versions of VP1 were obtained by removing the C-terminal end sequence exposed at the capsid surface of E11 (11-mer: PDTVKPDVSNH) and CVA9 (18-mer: TTVAQSRRRGDMSTLNTH). To abrogate the interfaces of Subtilisin A and BPT with CVA9 and CVB5 DE loops, the 14 amino acids of each loop (CVA9: QDPGTTLAQDMPVL, CVB5: EQSTIQGQDSPVL) were substituted by seven successive repeats of LV (LVLVLVLVLVLVLV). Targeted single substitution were further done on both DE loop and C-terminal end of CVA9 VP1, following a mass spectrometry identification of effective Subtilisin A cleavage sites on those two regions. Substitutions were chosen based on hydropathy property (H_i_) conservation, using the Kyte and Doolitle’s index^14^. As such, all T (T132, T282, T283) were individually substituted with a S (H_i T_ = -0.7, H_i S_ = - 0.8), Q135 with a N (H_i Q_ = H_i N_ = -3.5), L133 with a I (H_i L_= 3.8, H_i I_ = 4.5) and A134 with a V (H_i A_= 1.8, H_i V_ = 4.2). The effect of each modification on protease/virus interaction was assessed using AF2-M using the methodology described above.

### Docking refinement

The webserver HADDOCK 2.4^15^ was used to refine the interaction of Subtilisin A with each residue involved in an effective cleavage either in the DE loop (T132, L133, A134, Q135) or the C-terminal end VP1 sequence (T282, T283). For each analysis, both CVA9 protomer and Subtilisin A structures predicted with AF2-M were used as input. The VP1 was defined as protein chain A, and each targeted residues cited above was individually selected as active residue during the docking process. For Subtilisin A, defined as chain B, the four residues involved in the catalytic reaction (H32, D63, N154, S220) were selected as active residues. To assess the best conditions of analysis, we ran two different pipelines either adjusted to study protein/protein docking or peptide/protein docking. In the first experiment, all default settings were used for the analysis. In the peptide/protein docking, analysis parameters were adapted as follows: the number of structures for semi-flexible refinement, for final refinement and the number of structures to be analyzed were fixed at 400. The number of MD steps during it1 was set at 1000 steps on the high-temperature rigid body and during the first cooling stage of the rigid body, while this number during the second and third cooling stages was set at 3000.

For both pipelines, and regarding the viral residue targeted for the docking, DE loop and C-terminal end sequence flexibility were considered either automated, semi-flexible or fully flexible during the analysis. For all outputs, the Fraction of Common Contacts (FCC)^16^ method was used to filter the five best clusters returned by the analysis. As second filtering step, a distance analysis between the nucleophilic serine of Subtilisin A (S220) and each targeted viral residues was measured and a 3Å cut-off was applied as strong hydrogen bond indicator. Finally, the PRODIGY webserver was used to assess the overall portion of ICs.

### Proteins visuals, structure editing

All protein structure visuals, which include capsid viral proteins (protomers and viral assemblies), as well as enzyme monomers and enzyme/virus interactions, and inter-residue distance measurements were performed using ChimeraX (version 1.5)^17^. All visuals were then accommodated for publication using Adobe Illustrator (version 25.1).

### Data analysis

Data normality assessment and statistical analyses used to compare both viral infectivity and surface expression of integrins were performed with GraphPad Prism v.9.5.0. Samples comparison was done by one-tailed or two-tailed *t*-test depending on sample distribution, as precised in the caption of each corresponding figures. For the analyses showing a significant difference between samples variance (P < 0.05), a Welch’s correction was applied to each test. For all tests, an alpha value of 0.05 was used as a threshold for statistical significance.

### Data availability

SI videos can be found on the temporary Zenodo https://doi.org/10.5281/zenodo.8321541

## Acknowledgments

We thank EPFL collaborators for valuable discussions or technical support: L.F.K, M.DP, M.L. We thank the Flow Cytometry Core Facility (EPFL) and M.P from the Proteomic Core Facility (EPFL) for the use of the equipment and assistance. We thank S.B. and C.S-K (Finnish National Institute for Health and Welfare) for providing the environmental CVA9 isolate. This research was funded by the Swiss National Science Foundation (grant no. 310003A-182468).

## Author contributions

Conceptualization: MHC, TK, BR

Methodology: MHC, SCD

Investigation: DC, MHC, SCD, BR, ST

Visualization: MHC

Data curation: MHC

Formal analysis: MHC

Project administration: MHC

Funding acquisition: TK

Supervision: MHC, TK

Writing—original draft: MHC

Writing—review & editing: MHC, SCD, TK

All authors read and approved the final version of the manuscript.

## Competing interests

Authors declare that they have no competing interests.

---

**Extended Data Fig. 1:**
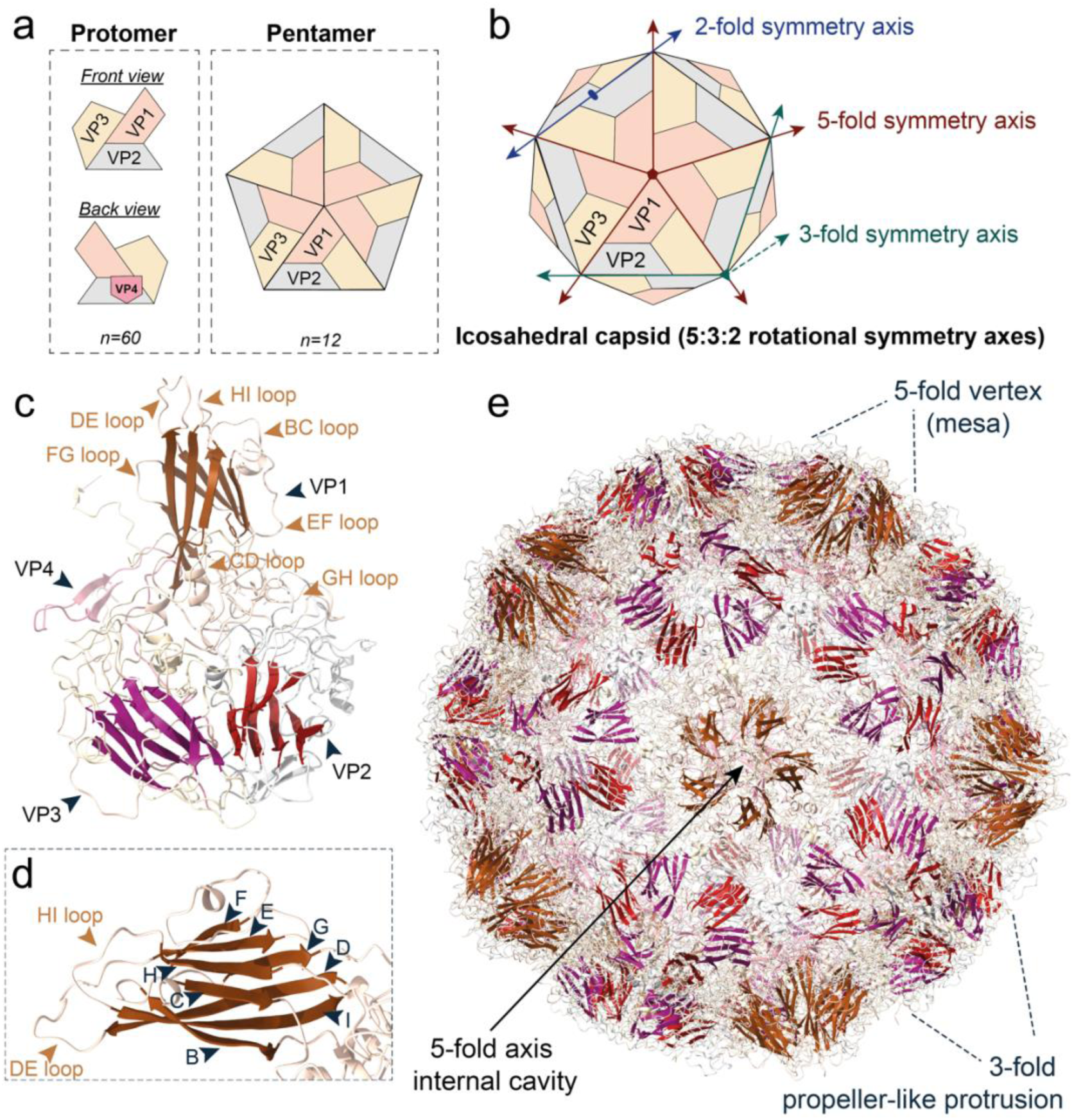
Main features of enterovirus capsid architecture. **a.** Description of one (protomer) and five (pentamer) viral subunit(s) composing the capsid of enteroviruses. **b.** Diagram of the icosahedral capsid shell describing a 5:3:2 rotational symmetry axis. **c.** Ribbon representation of CVA9 protomer (PDB:1d4m), displaying three jelly rolls folds. The seven loops interconnecting the jelly roll fold segments of VP1 are shown in orange. **d.** Ribbon representation of a magnified view of the VP1 jelly roll fold ß-sandwich, indicating a protein segments nomenclature from B to I. **e.** Ribbon view of CVA9 full capsid structure showing the protein protrusions induced by the assembly of viral proteins on the symmetry axes centers, and the open circular cavity of the 5-fold axis. *(Figure based on published data: see references in the Introduction section)*.

**Extended Data Fig. 2:**
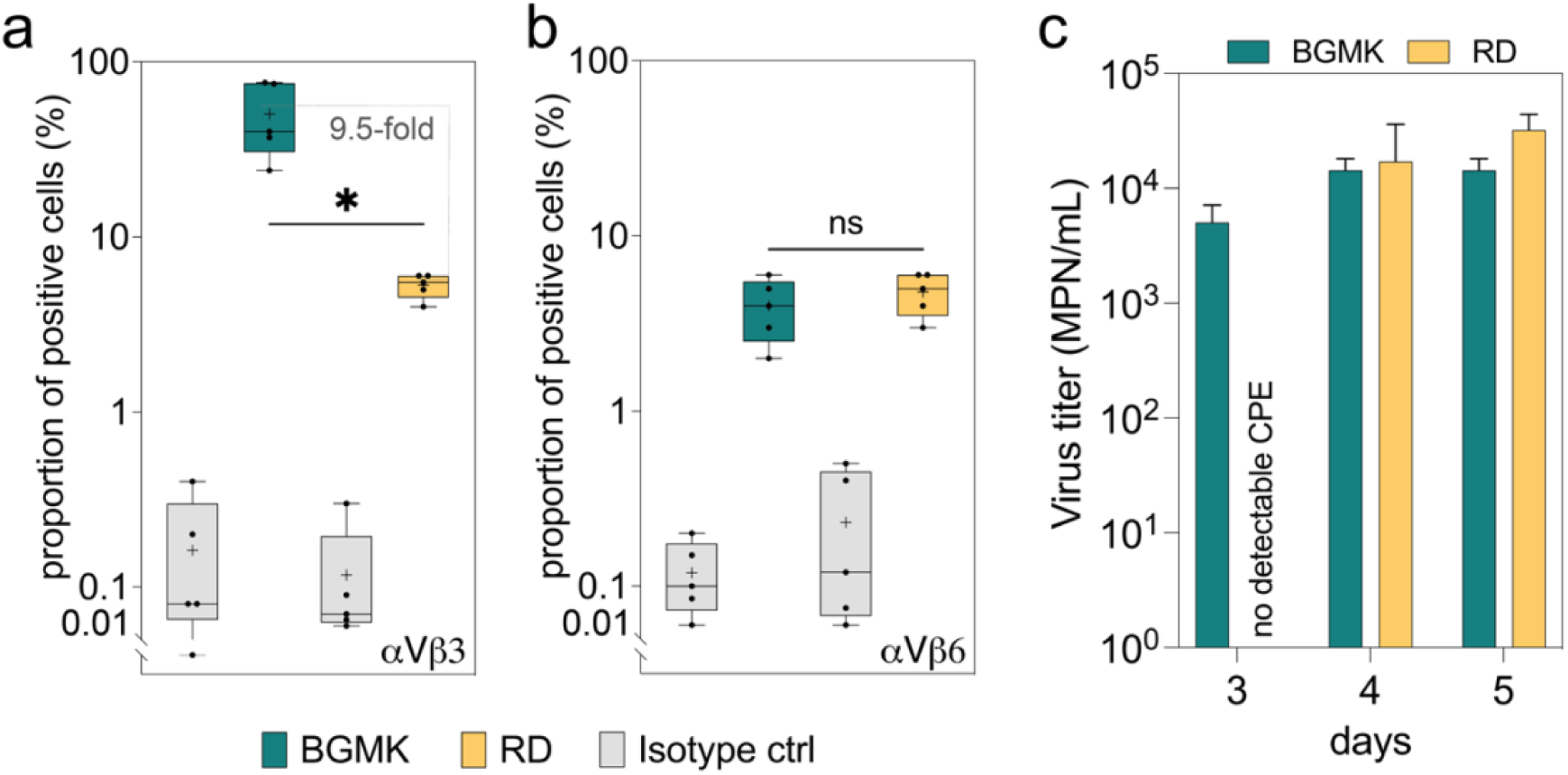
Infectivity properties of CVA9 on BGMK and RD cells. **a-b.** Flow cytometry analysis of the percentage of BGMK and RD cells expressing **(b)** αVβ3 (t=4.254, p-value:0.0130 (*); two-tailed unpaired t-test, n=5) and **(c)** αVβ6 (t=0.873, p-value: 0.4091 (ns); two-tailed unpaired t-test, n=5) at the cell surface. For each box plot, the horizontal bar within the interquartile range corresponds to the median of values and ‘+’ indicates the mean. The upper and lower limits of dataset distribution correspond to minimum and maximum values. **c.** CVA9 infectivity monitoring on RD cells and BGMK cells 3, 4 and 5-days post-infection.

**Extended Data Fig. 3:**
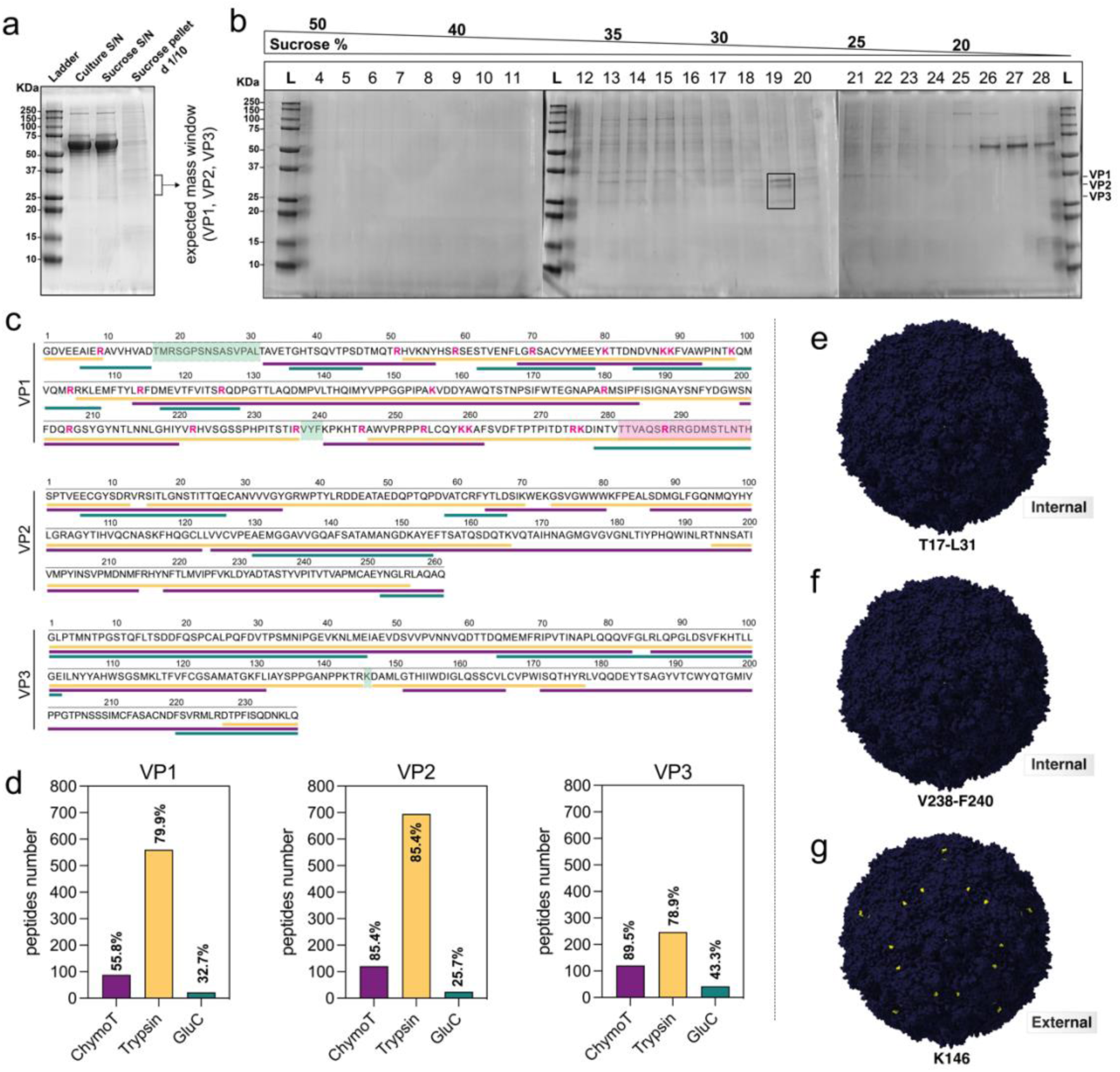
Mass spectrometry identification of CVA9 (environmental isolate) viral protein sequences following a two-step sucrose virus purification procedure. **a.** Protein profile analysis of CVA9 pre-purification on 20% sucrose cushion. Culture S/N corresponds to cell culture supernatant containing native virions. Sucrose S/N corresponds to sucrose supernatant. Sucrose pellet d_1/10_ refers to the native CVA9 pellet enriched by the cushion and diluted to 1/10 for protein gel analysis. **b.** Protein profile analysis of CVA9 purification on discontinuous sucrose gradient. Each number indicates the order of the 500 μL fractions extracted from the bottom of the ultracentrifugation tube. The black box (fraction 19) indicates the purified viral capsid proteins VPs (*i.e*., VP1, VP2 and VP3). **c.** Protein sequences of CVA9 following in-gel digestion of each VP by chymotrypsin, trypsin, or GluC prior to LC-MS analysis. Each colored bar under sequences represents the sequence coverage of the working strain on the reference strain CVA9 Griggs (extracted from PDB:1d4m), resulting from each enzymatic digestion: yellow (trypsin), purple (chymotrypsin), green (GluC). Pink letters in the VP1 sequence correspond to amino acid residues for which a cleavage of the C-terminal peptide bond was observed after trypsin digestion. The light green boxes correspond to the sequences non-covered by the analysis. The pink box corresponds to the C-terminal end of VP1 carrying the RGD motif. **d.** Number of unique peptides resulting from the enzymatic digestion for each individual VP and associated coverage percentage. **e-g.** Surface representation of a full capsid particle of CVA9 (strain Griggs, 1d4m), highlighting the internal or external position of the 19 residues not resolved by LC-MS. The K146 of VP3, is the only external residue for which the identity remains unclear in CVA9 sequence. Each call of residues on the surface capsid has been done using ChimeraX. Capsids visuals have been oriented on the 5-fold axis center.

**Extended Data Fig. 4:**
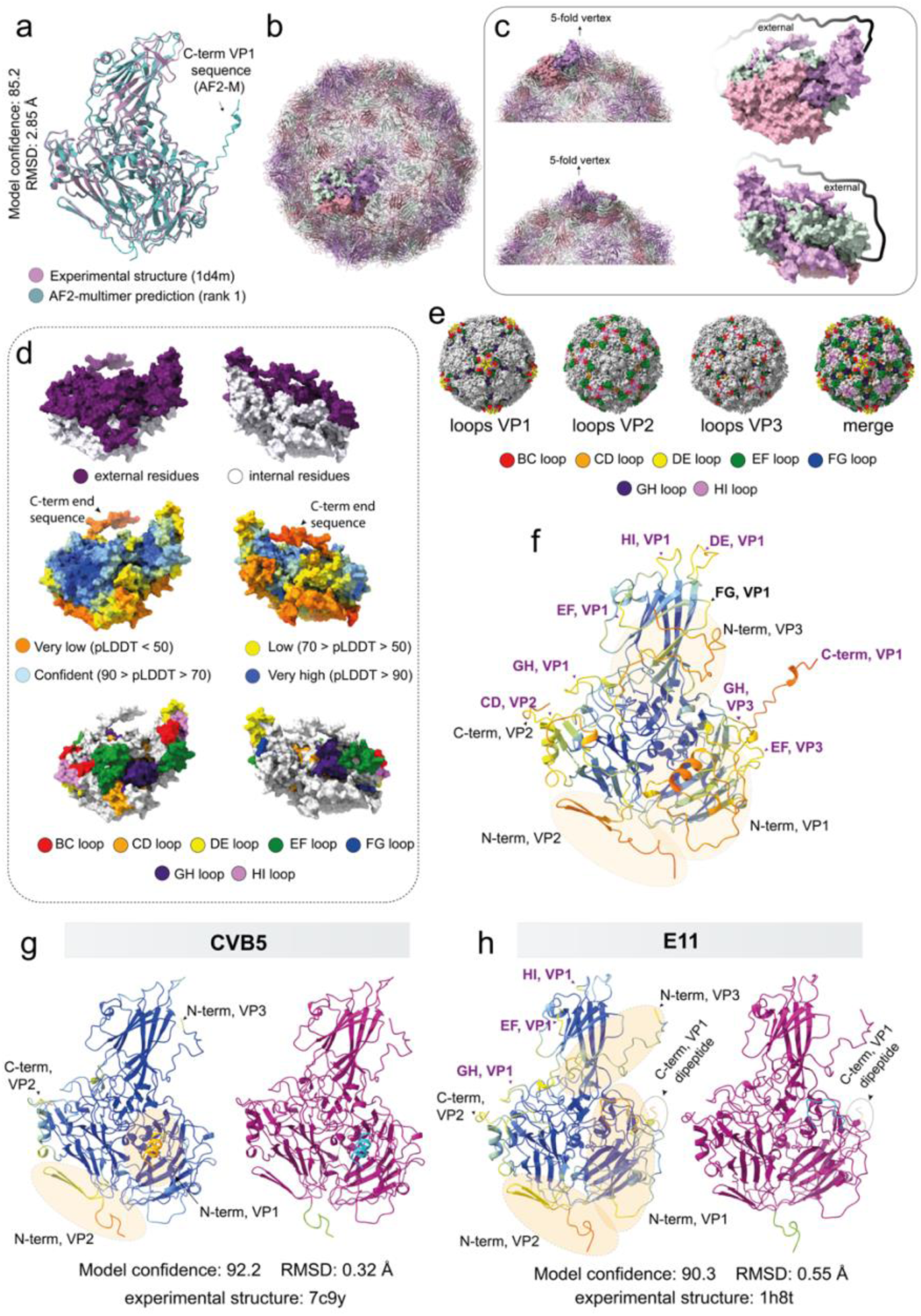
Additional visuals of *Enterovirus* protomers folding using AF2-M. **a.** Overlay of the experimental structure (1d4m) with the modeled structure (rank 1) of CVA9 protomer. **b.** Surface view of a CVA9 protomer embedded on a capsid shell (ribbon view). **c.** Left view (top left) and right view (bottom left) of a CVA9 protomer embedded on a capsid shell and associated protomer modeled (rank 1) oriented on the same direction (top and bottom right). **d.** Surface view of a modeled CVA9 protomer (rank 1) oriented on the capsid shell as shown in **c.** From top to bottom, color scales correspond to the exposure of residues on the outer or inner side of the capsid (purple/white), the confidence of folding performed with AF2-M (plDDT scale) and the exposure of jelly roll fold loops for each VP of the protomer (rainbow scale color). **e.** Surface view of CVA9 capsid showing the exposure of jelly roll folds associated loops for each VP. The “merge” surface view is based on the sum of all loop residues exposed for all VPs. **f.** Ribbon diagram of CVA9 protomer modeled by AF2-M (rank 1), displaying the confidence level of the model colored by plDDT. The eight areas noted in purple correspond to protein regions exposed on the capsid surface and predicted with low confidence (plDDT < 70). Other annotations in black correspond to regions predicted with low confidence but not exposed on the external capsid surface. **g-h.** Ribbon diagrams of CVB5 **(g)** and E11 **(h)** modeled using AF2-M (rank 1), displaying on the left the confidence level of the model colored by plDDT and on the right the confidence according to PAE score matrix. On each model colored by plDDT, all areas pointed by an arrow correspond to protein regions predicted with low confidence. On **h**, the circled area corresponds to the additional dipeptide found at the C-terminal end of VP1 (external segment) which was absent from the experimental structure 1h8t but modeled by AF2-M.

**Extended Data Fig. 5:**
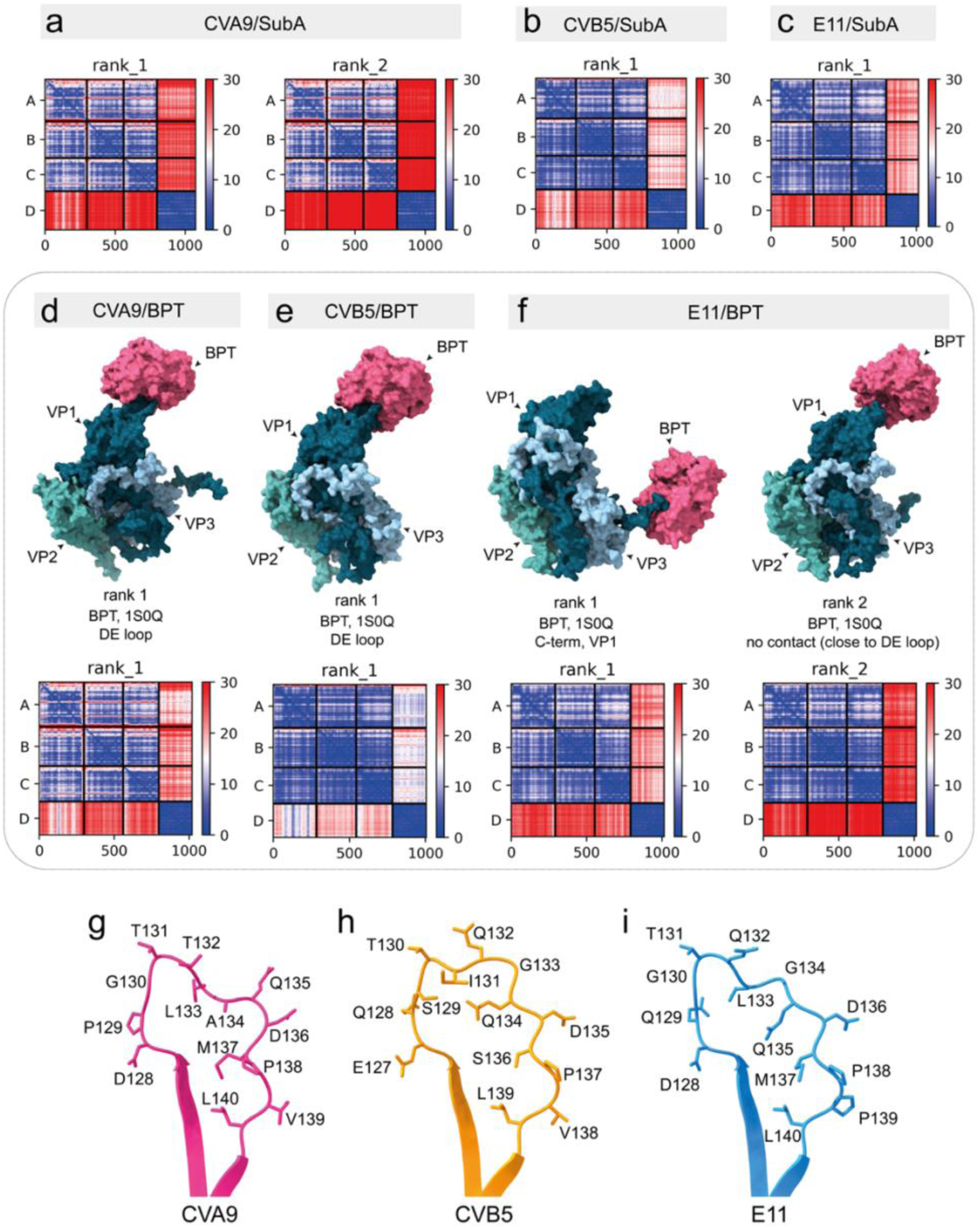
Additional data showing predictive binding of BPT to the DE loop and/or to the C-terminal end VP1 sequence of enteroviruses. **a.** PAE plots of the two best rank models resulting from the prediction of interfaces between Subtilisin A and CVA9 protomer with AF2-M. **b.** PAE plot of the best rank model of CVB5/Subtilisin A interface predicted by AF2-M. **c.** PAE plot of the best rank model of E11 /Subtilisin A interface predicted by AF2-M. **d.** Surface view of the best model of interaction between BPT and CVA9 protomer returned by AF2-M and associated PAE plot. **e.** Surface view of the best model of interaction between BPT and CVB5 protomer returned by AF2-M and associated PAE plot. **f.** Surface view of the two best rank models of interaction between BPT and E11 protomer returned by AF2-M and associated PAE plots. **a-f.** For each PAE plot, letters A, B, C and D correspond to VP1, VP2, VP3 and the protease of interest, respectively. **g-i.** Magnified view of the ribbon representation of the D and E segments of VP1 from (**g**) CVA9, (**h**) CVB5 and (**i**) E11 and amino acid composition of the DE loop, showing the absence of any basic residue (R, K).

**Extended Data Fig. 6:**
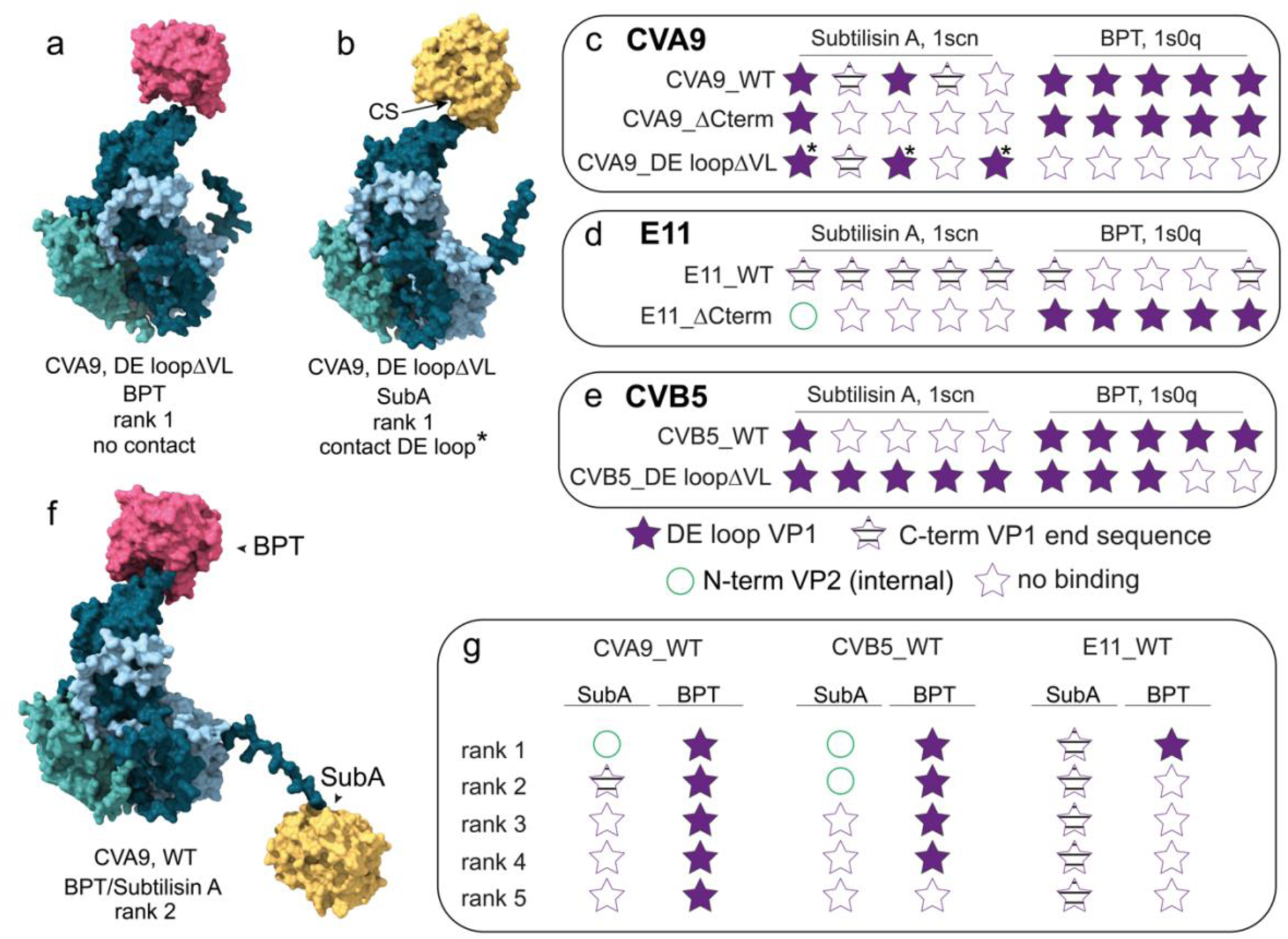
Protease/capsid interfaces are virus-and protease-specific. **a.** Surface view of BPT recruitment on CVA9 protomer showing the loss of interface between the viral DE loop and the protease after the full substitution of the DE loop by a VL repeat sequence. **b.** Surface view of Subtilisin A recruitment on CVA9 protomer a partial loss of interface between the viral DE loop and the protease after the full substitution of the DE loop by a VL repeat sequence. **c.** Overview of predicted interfaces of Subtilisin A and BPT with CVA9 protomer (CVA9_WT), compared with the protomer version whose C-terminal sequence has been completely truncated (CVA9ΔCterm) or with the hydrophobic DE loop protomer mutant (CVA9_DE loopΔVL). **d.** Overview of predicted interfaces of Subtilisin A and BPT with the E11 protomer (E11_WT), compared with the protomer version whose C-terminal sequence has been completely truncated (E11ΔCterm). **e.** Overview of predicted interfaces of Subtilisin A and BPT with CVB5 protomer (CVB5_WT), compared with the hydrophobic DE loop protomer mutant (CVB5_DE loopΔVL). **f.** Surface view of a simultaneous recruitment of BPT and Subtilisin A to both CVA9 protomer IDPs. **g.** Overview of the predictive interfaces during a modeling in presence of the two serine proteases for each viral protomer. (CS) catalytic site, (*) used to indicate predictive outputs showing an interaction between Subtilisin A and the DE loop, for which the CS is no longer aligned with the viral sequence. All AF2-M predictions have been done using 3 recycles.

**Extended Data Fig. 7:**
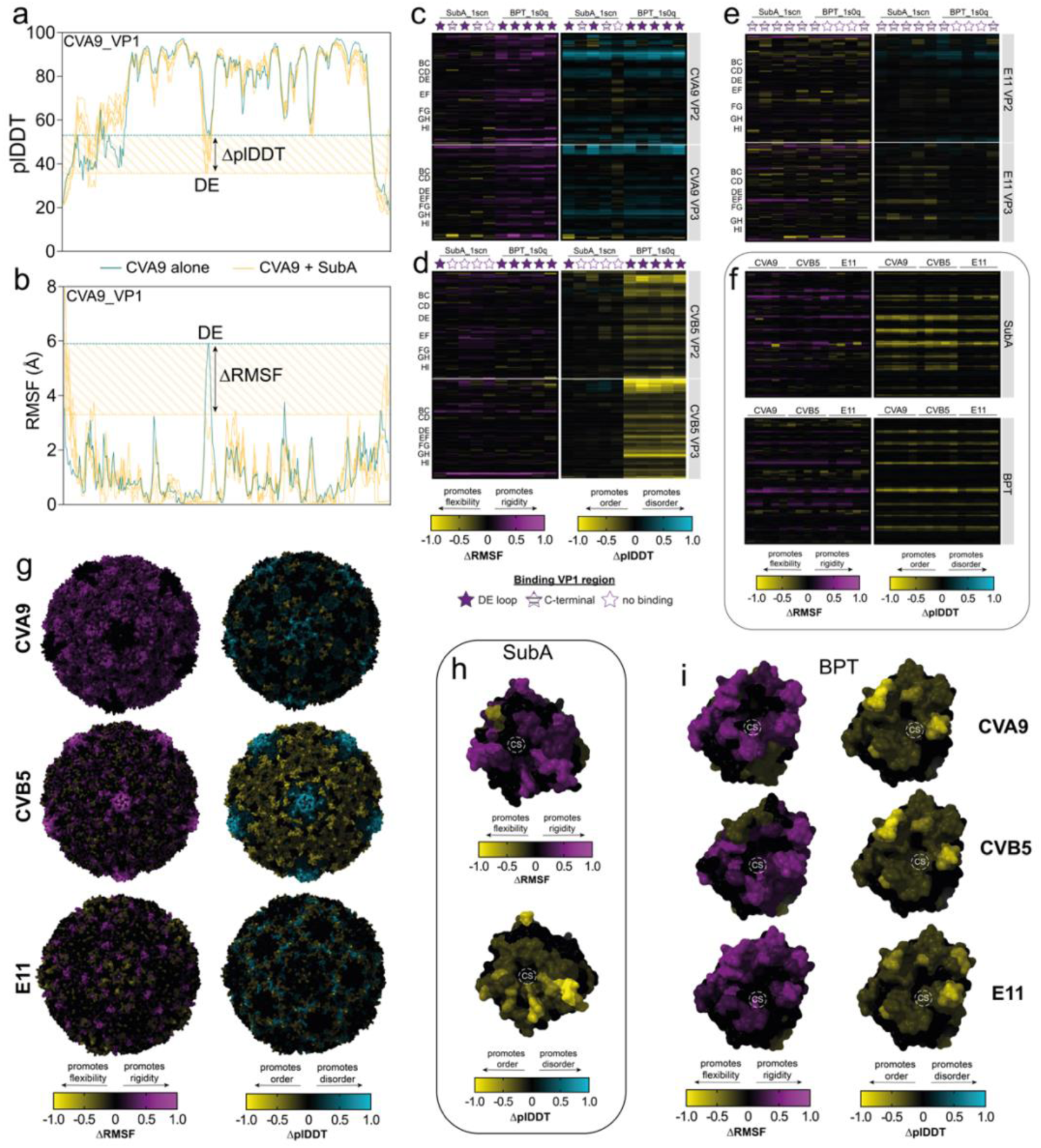
Additional data regarding both viral proteins and proteases plasticity upon the computational exposure of enzymes to enteroviruses protomers. **a.** Overlay of plDDT profiles of CVA9 VP1 achieved without exposure to Subtilisin A (green) or with the protease (yellow). The striped area indicates the plDDT variation (ΔplDDT) measured for the DE loop in response of Subtilisin A presence in the computational field. **b.** Overlay of RMSF profiles of CVA9 VP1 measured without exposure to Subtilisin A (green) or with the protease (yellow). The striped area indicates the RMSF variation (ΔRMSF) measured for the DE loop in response of Subtilisin A presence in the computational field. **c-e.** Distribution of VP2 and VP3 flexibility (ΔRMSF) or disorder (ΔplDDT) variations induced by Subtilisin A or BPT for (c) CVA9 (d) CVB5 and (e) E11. On the top of each heatmap, stars indicate the predictive binding regions for each protease used, as mentioned above. On the top end of each heatmap is indicated the name of the protease used in the analysis. For each heatmap, the upper part corresponds to VP2, and the bottom part to the VP3. **f.** Distribution of Subtilisin A (top) and BPT (bottom) flexibility (ΔRMSF) or disorder (ΔplDDT) variations induced by an exposure with each virus type. **g.** Surface view of each viral capsid colored by ΔRMSF (left part) or by ΔplDDT (right part) for each virus type following an exposure with BPT. All capsids have been centered on the 5-fold axis view. **h.** Surface view of Subtilisin A colored by ΔRMSF (left part) or by ΔplDDT (right part), after an exposure with CVB5 protomer. **i.** Surface view of BPT colored by ΔRMSF (left part) or by ΔplDDT (right part), after an exposure with CVA9, CVB5 or E11. (CS): catalytic site. All profiles variations have been calculated based on AF2-M models folded using 3 recycles.

**Extended Data Fig. 8:**
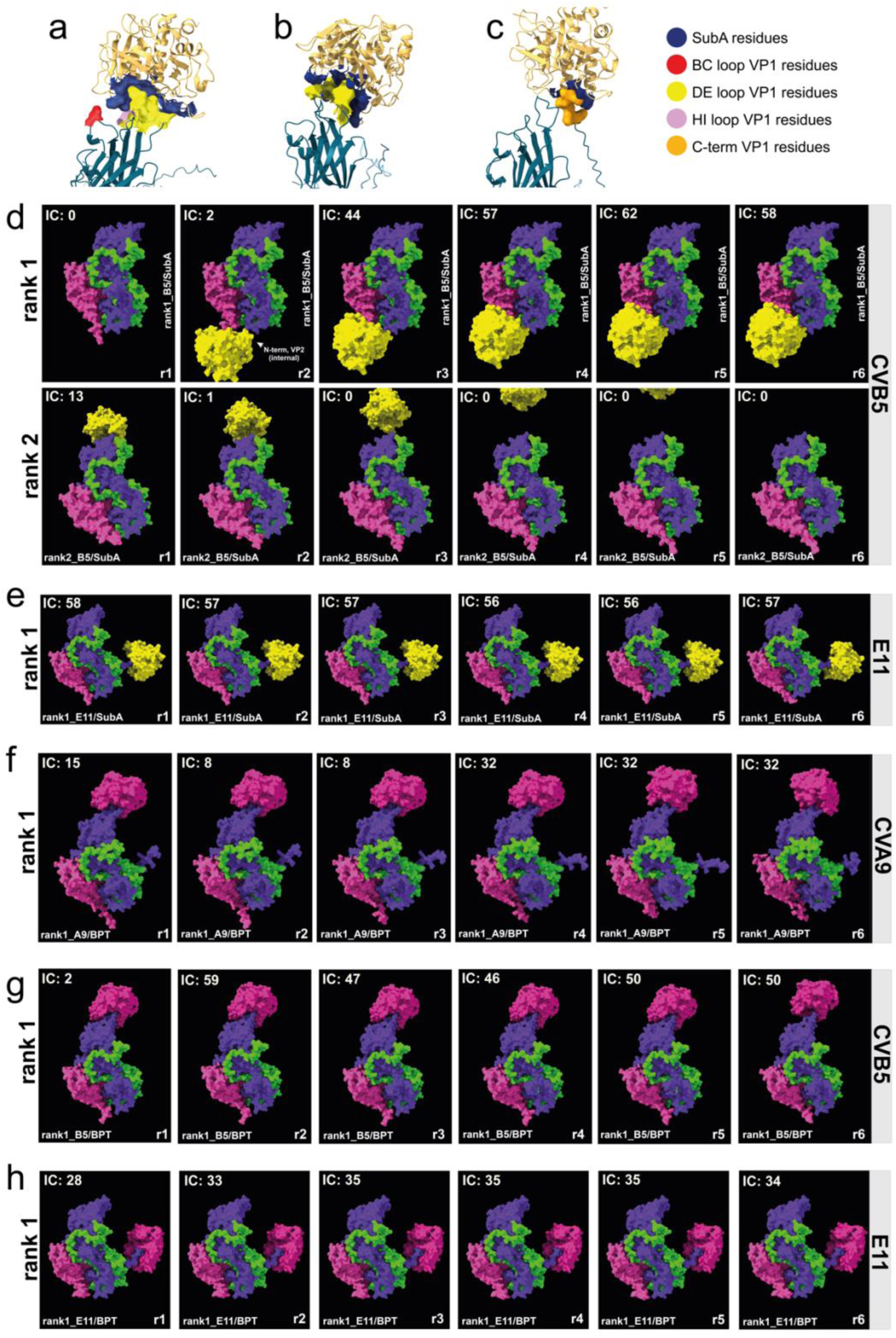
Additional predictive visuals and timelines of Subtilisin A and BPT recruitments on CVA9, CVB5 and E11 protomers. **a.** Surface area contact between Subtilisin A and CVA9 5-fold vertex captured after the second recycle (rank1, r2, IC_max_ = 54). **b.** Surface area contact between Subtilisin A and CVA9 5-fold vertex captured after the second recycle (rank2, r2, IC_max_ = 30). **c.** Surface area contact between Subtilisin A and CVA9 C-terminal end sequence captured after the fifth recycle (rank2, r5, ICmax = 2). Predictive Subtilisin A recruitment on (**d**) CVB5 and (**e**) E11. f-h. Predictive BPT recruitment on (**f**) CVA9, (**g**) CVB5 and (**h**) E11. Each timeline is composed of 6 recycles and the number of interatomic contacts (IC) is specified in each frame.

**Extended data Fig. 9:**
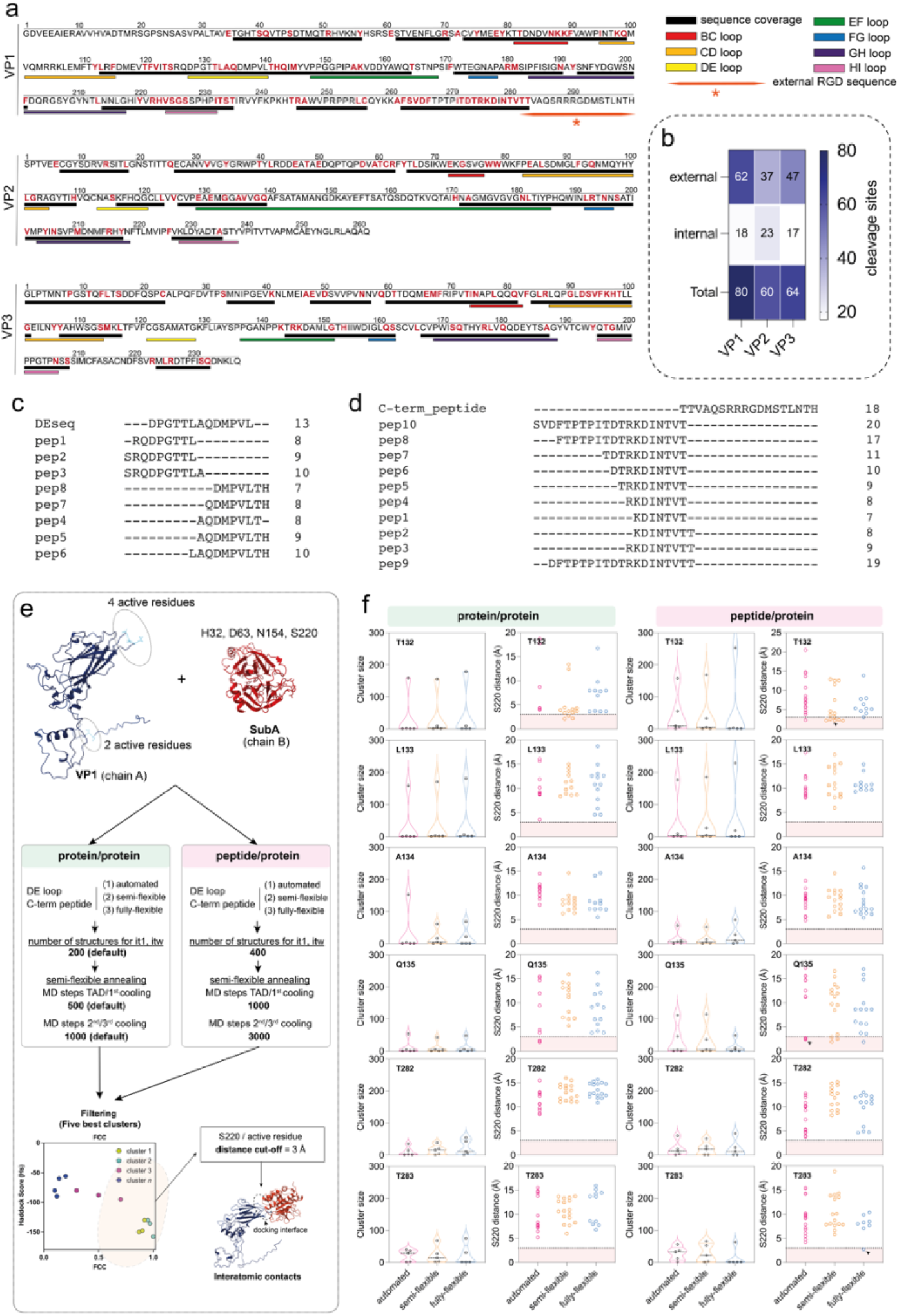
Additional data of the deductive inference process used to identify VP1 residues involved in capsid/Subtilisin A interaction leading to viral inactivation. **a.** Subtilisin A cleavage points on CVA9 VP1, VP2 and VP3. Red letters correspond to amino acid residue for which a cleavage of the C-terminal peptide bond was observed. The different colored areas below the sequences correspond to the sequence coverage obtained following in-gel digestion of VP with Subtilisin A or to the different areas of interest in this work and are explained in the legend on the right. **b.** Number and structural distribution of unique cleavage sites identified on each VP following 30 minutes of in-gel protein digestion with Subtilisin A. **c.** Full list of unique peptides from the DE loop identified by mass spectrometry after in-gel digestion of VP1 protein with Subtilisin A. **d.** Full list of unique peptides of the region preceding the C-terminal end sequence of VP1 identified by mass spectrometry after in-gel digestion of VP1 protein with Subtilisin A. **e.** Directed-site docking procedure used to screen and prove the docking of Subtilisin A on biologically cleaved residues. For each active viral residue, the procedure was run by either adapting or not the flexibility of each protein portion of interest, and by adjusting or not certain analysis parameters during the iterations. The five best clusters were selected using a fraction of common contact (FCC) filtering and the results were screened by applying a cut-off distance of 3Å between the nucleophilic serine of Subtilisin A (S220) and each selected active virus residue. **f.** Distribution of the sizes of the five best clusters for each procedure used (violin plot, left) and distribution of the distances between the S220 and the selected viral residue for each procedure (scatter plot, right). The colored areas below 3Å correspond to the distances selected to screen the relevant docking structures. Each arrow in the scatter plots from the protein/peptide docking corresponds to the three models selected for Figure 5.f.

**Extended data Fig. 10:**
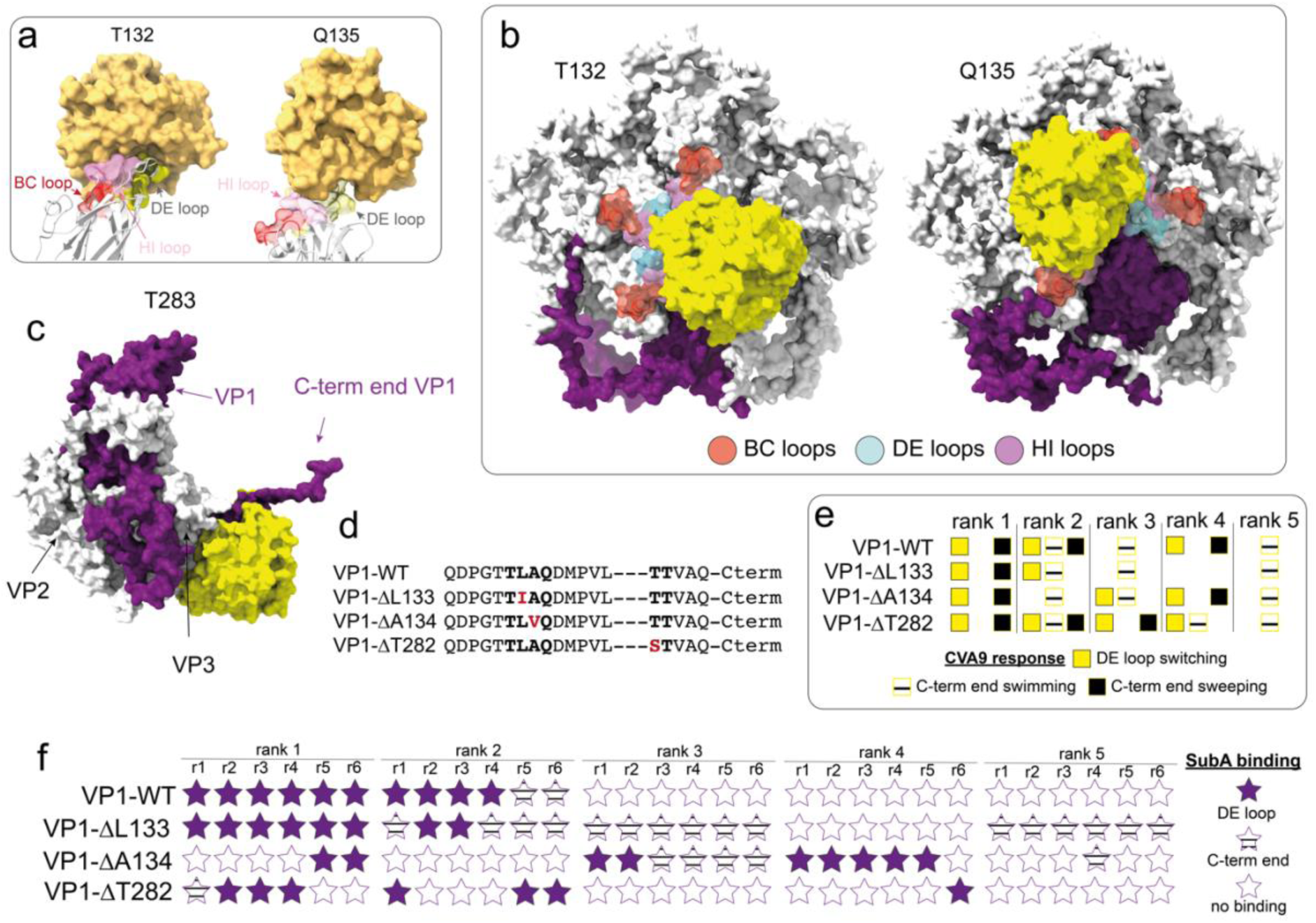
Additional visuals of VP1/Subtilisin A docking and discreditation of the last three structurally inaccessible amino acids in Subtilisin A recruitment to CVA9 capsid. **a.** Surface representation of Subtilisin A docked to the DE loop of VP1 on residues T132 and Q135. For residue T132, the BC, DE and HI loops contribute to the virus/capsid interface, while for residue Q135, only the DE and HI loops participate to the interface. **b.** Subtilisin A docked to the VP1 DE loop on residues T132 and Q135, fitted to a VP1 pentamer. **c.** Docking of Subtilisin A on VP1 C-terminal end T283 residue. The VP1/Subtilisin A docking structure obtained with Haddock 2.4 (purple/yellow) and the modeled protomer (AF2-M, rank1) of CVA9 (white) were aligned on VP1 with ChimeraX. The result indicates that the Subtilisin A docked to VP1 also locally fits on VP3 surface (External part of the capsid). **c.** Additional *in-silico* point mutations of VP1 L133, A134 and T282 residues cleaved by Subtilisin A during a in-gel digestion of VP1 (denatured VP1) but unconfirmed by structural analysis. On the left of the dotted line, the sequence corresponds to the DE loop and on the right, the sequence corresponds to the five first amino acid of the C-terminal end of VP1. **d.** Influence of CVA9 VP1 point mutations on the prediction of DE loop and C-term end trajectories during Subtilisin A recruitment to viral capsid. The absence of a square indicates that the corresponding trajectory event was not observed in any recycle of the analysis. **e.** Influence of CVA9 VP1 point mutations on the ability to Subtilisin A to bind the DE loop or the C-terminal end sequence. Subtilisin A recruitments on CVA9 protomer were predicted with AF2-M using 6 recycles.

## Notes

https://doi.org/10.5281/zenodo.8321541

### Competing Interest Statement

The authors have declared no competing interest.

### Summary of Updates

SI videos are available at https://doi.org/10.5281/zenodo.8321541

